# A deformable attractor manifold organizes human resting-state brain dynamics

**DOI:** 10.64898/2025.12.02.691788

**Authors:** Anastasios P. Athanasiadis, Marmaduke M. Woodman, Meysam Hashemi, Paul Triebkorn, Spase Petkoski, Viktor K. Jirsa

## Abstract

Intrinsic brain activity is often described as wandering within a continuous multivariate space, yet the organizing principles that constrain these dynamics remain unclear. Here, we show that spontaneous human brain activity during rest is structured by a deformable attractor manifold. Using large-scale fMRI datasets and a latent dynamical model, we find that cortical activity occupies two reproducible regimes: a low-coherence state with a unimodal latent distribution and a high-coherence state that exhibits bimodality, consistent with transient bistability across association networks. A compact two-parameter energy landscape explains these dynamics, revealing that transitions arise not from switching between discrete states, but from continuous deformation of the manifold that reshapes attractor geometry. Excursions into the bistable regime occur as rapid “jumps”, whereas returns follow slow drifts along the manifold, reflecting network-specific timescales. Individuals with greater expression of the bistable regime show higher cognitive fluidity, and manifold parameters differentiate mild cognitive impairment from matched controls. These findings identify an organizing geometric and dynamical principle of resting activity, linking large-scale cortical coordination, cognitive variability, and vulnerability to pathology.

## Introduction

Unlike non-living matter such as ferromagnets, which require external influence to align their domains of atomic spins and become magnets, the brain, as a complex hierarchical system, possesses the remarkable ability to self-organize (Haken, 1977; Salthe, 1985; Kelso, 1995). This inherent property gives rise to phenomena like sleep and functional spatio-temporal patterns of activity observed during rest (Buzsáki and Draguhn, 2004), known as resting-state networks (RSNs) (Biswal et al., 1995; Thomas Yeo et al., 2011; Power et al., 2011). Hierarchy is spatial, as brain activity originates from the fundamental excitability of individual neurons and constrained by the intricate anatomy and geometry of the cerebral cortex (Atasoy et al., 2016; Pang et al., 2023; Vohryzek et al., 2025); and temporal, as higher cognitive areas that are responsible for integrating complex information, typically operate on longer timescales than sensory/motor areas (Kiebel et al., 2008; Honey et al., 2012; Murray et al., 2014). The produced dynamics ultimately fosters a delicate balance between localized and distributed processing across diverse brain areas and networks (Damasio and Damasio, 1994; Ito et al., 2020), which translates into the observed alternating synchronization and desynchronization of different networks during rest (Honey et al., 2007; Deco et al., 2011; Raichle, 2011).

In resting-state functional magnetic resonance imaging (fMRI), this ongoing reconfiguration of functional networks generates a variety of blood oxygenated level-dependent activation (BOLD) patterns, known as coactivation patterns. This intrinsic temporal variability led to the development of functional connectivity dynamics (FCD) analysis methods (Hutchison et al., 2013; Hansen et al., 2015), that can capture how functional dynamics shifts between epochs of global and modularized (anti-)correlations (Zalesky et al., 2014; Rabuffo et al., 2025). Mathematically, coupled non-linear dynamics provides a natural language for formulating such coordination dynamics and self-organization in terms of instabilities, timescales separation, and dimensionality reduction (Haken, 1977; Schöner and Kelso, 1988; Kelso, 1995). In this regard, macroscopic brain patterns are frequently viewed as emergent functional modes or states, representing coordinative structures or synergies (Hashemi et al., 2025). The natural habitat of these modes is a low-dimensional state space or *manifold*, and the time evolution of their probability density dynamics is captured from the Fokker-Planck equation (Risken, 1989; Friston et al., 2023).

Numerous studies have demonstrated that task-based activity across different species and modalities can be represented by flows using low-dimensional embeddings (operationalized by Schneider et al. (2023)), including human fMRI signal (Shine et al., 2019). Such processes can be described by *structured flows on manifolds* (SFMs), a dynamical framework that builds upon self-organization, where flows become the focus that shapes coordinated transitions rather than fixed modes (attractors) (Huys et al., 2014; Pillai and Jirsa, 2017). However, co-activation patterns at rest have been treated as stationary, i.e. time-invariant attractor states, as whole-brain modeling has been employed to characterize the resting-state manifold (Deco et al., 2012; Hansen et al., 2015; Golos et al., 2015; Englert et al., 2024). Stationary attractors are either fixed points or (approximately) continuous (e.g. line, ring attractor) (Khona and Fiete, 2022) and reflect abstract representations, encoding a multitude of possible patterns in a compressed form (Ghosh et al., 2008; Hansen et al., 2015). Empirically, mere classification of such patterns has already yielded significant insights into the nature of the activity, revealing aspects such as non-stationarity (Liu and Duyn, 2013), individual “fingerprinting” capabilities (Peng et al., 2023), and relevance to information processing in cognitive tasks (Petersen and Sporns, 2015). Significantly, the latter has been linked to the hypothesis that spontaneous activity is a signature of the brain’s internal model and priors which interact with task-evoked patterns (Raichle, 2011; Petersen and Sporns, 2015; Dehaene and Cohen, 2007). However, evidence about how the resting-state manifold is organized to give rise to this range of configurations remains elusive (Fousek et al., 2024).

Co-(de)activation events (CAEs) at rest denote bursty transients generated by the strongest (thus rare and global) co-(de)activations of brain areas across time (Esfahlani et al., 2020; Rabuffo et al., 2025). When projected onto the manifold, trajectories that exhibit CAEs form distinct structured flows separable from the core activity (Fousek et al., 2024). The “labile brain” hypothesis (Friston, 2000) highlighted the crucial role of short-lived transients, describing how brain dynamics move from a state of *stable incoherence* through *dynamic instability* to *complete entrainment*. This work expands beyond the CAEs or co-activation patterns (Iraji et al., 2022), and focuses on stationarity to characterize the resting-state manifold. Jirsa and Sheheitli (2022) showed how the stationary Fokker-Planck density function gives access to the underlying SFMs and links to the free energy principle (Friston, 2013; Friston et al., 2023). Building upon findings that demonstrated natural partitioning of time-varying functional connectivity based on coherence (Zalesky et al., 2014; Kong et al., 2021), we aim to characterize the stationary modes and flows during two distinct states of dynamics at rest: a low coherence state (LCS) marked by widespread incoherence, and a high coherence state (HCS) exhibiting overall coordinated activity. We describe their temporal and whole-brain manifold organization, and examine the geometries of the RSNs’ potential functions (also viewed as the information-theoretic *surprise* (Friston et al., 2023)) to understand the composition of LCS and HCS. Subsequently, we reveal the attractive flow on the manifold for both states. Finally, to analyze state transitions, we assess non-stationarity and the role of CAEs in shaping them.

## Results

### Identification of stable LCS and transitory HCS of activity links to *fluidity*

Empirical fMRI data (repetition time (TR)=0.72s) from the Human Connectome Project (HCP) was used to examine BOLD activity from (P=200) unrelated participants during rest. To ensure reproducibility, we repeated our analysis on the second day (day-2) data set and a high-resolution subset (TR=0.6) of the Alzheimer’s Disease Neuroimaging Initiative (ADNI) dataset, which consisted of elderly control (CN) (P=18), mildly cognitive impaired (MCI) (P=23) and Alzheimer’s disease (AD) patients (P=7) (Methods A). For each participant (p) we extracted the instantaneous functional connectivity (FC) of the phases of brain areas (Cabral et al., 2017) (Fig. 1A). Averaging the overall links yielded the global (mean) coherence, which was used to obtain the LCS and HCS (details in Methods B) (Fig. 1B). As expected, the HCS signified epochs of reduced dimensionality and more coordinated activity than the LCS, quantified by the spectral radius of the FC matrices (Fig. S1) (Hancock et al., 2024). Additionally, the correlation of the FC matrices across time yielded the FCD matrix per participant that demonstrates the recurrent occurrence of brain coactivation patterns (Hansen et al., 2015) (Fig. 1D left). The variance of the FCD matrix is linked to a metric of brain fluidity, which has been associated with cognitive flexibility (Senden et al., 2017; Naik et al., 2017), different consciousness levels (Breyton et al., 2023), healthy aging (Lavanga et al., 2023) and tau-protein related characterization of AD patients (Mazzara et al., 2025).

**Figure 1.**
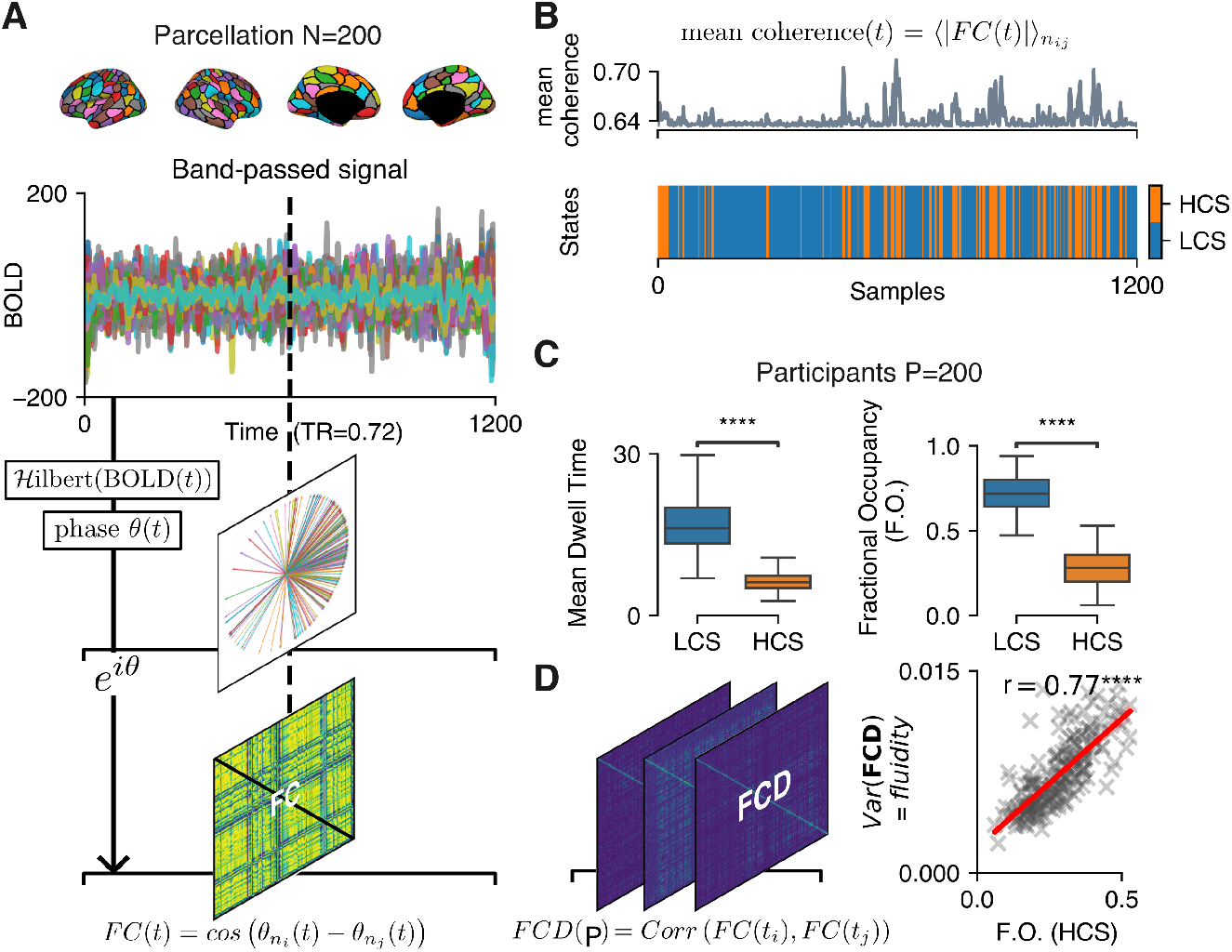
Different signal coherence levels at rest identify temporally stable Low Coherent State (LCS) and short-lived High Coherent State (HCS) of activity that is linked to *fluidity*. (**A**) Parcellated BOLD signal was analyzed from resting-state fMRI: first, bandpass filtered between [0.01, 0.18] Hz, then transformed to the Hilbert space to extract the phase of each brain area. The cosine of the pairwise phase difference between all nodes at each time point yielded the instantaneous functional connectivity (FC) matrices. (**B**) The mean coherence of the signal was calculated by the mean absolute values of each FC matrix and was fitted by a hidden Markov model to predict the sequence of LCS and HCS in the signal. (**C**) Mean dwell time and fractional occupancy (FO) across all participants revealed the transient nature of HCS contrasting the temporal stability of LCS (*p* < 10^−4^). (**D**) The functional connectivity dynamics (FCD) was estimated by the correlation of all FC matrices for each participant p. The FCD variance has been linked with brain *fluidity*, and significantly correlated with the FO of the HCS state.

We calculated the proportion of time spent in each state, or fractional occupancy, and the average duration of the occupancy of each state, or mean dwell time, to characterize the states temporally. The significantly higher fractional occupancy (*p* < 10^−4^) and mean dwell times (*p* < 10^−4^) in LCS suggested a temporally stable nature for LCS, contrasting the transient HCS (Fig. 1C). The fractional occupancy of HCS significantly correlated with fluidity (*r* = 0.77, *p* < 10^−4^) (Fig. 1D right), implying that its temporal instability shapes the variability in FCD. This result was replicated in the two other data sets (Fig. S2B, C).

### HCS bimodality contrasts LCS unimodality and translates into richer variety of coactivation patterns

Next, we examined how the two states are represented in the resting-state manifold by applying principal components analysis (PCA) on the overall activity. For a test participant, we observed that the HCS activity was strongly dispersed across the first principal component (PC) compared to LCS (Fig. 2A top and C bottom). We aimed to examine this difference through the underlying stationary probability density functions (*f*_*st*_); thus, we identified non-stationary transitional events (Methods C). This yielded two types of non-stationarities; the fast and CAEs (Fig. 2A bottom), where both fell significantly more into the HCS across all participants (*p* < 10^−4^) (Fig. 2B).

**Figure 2.**
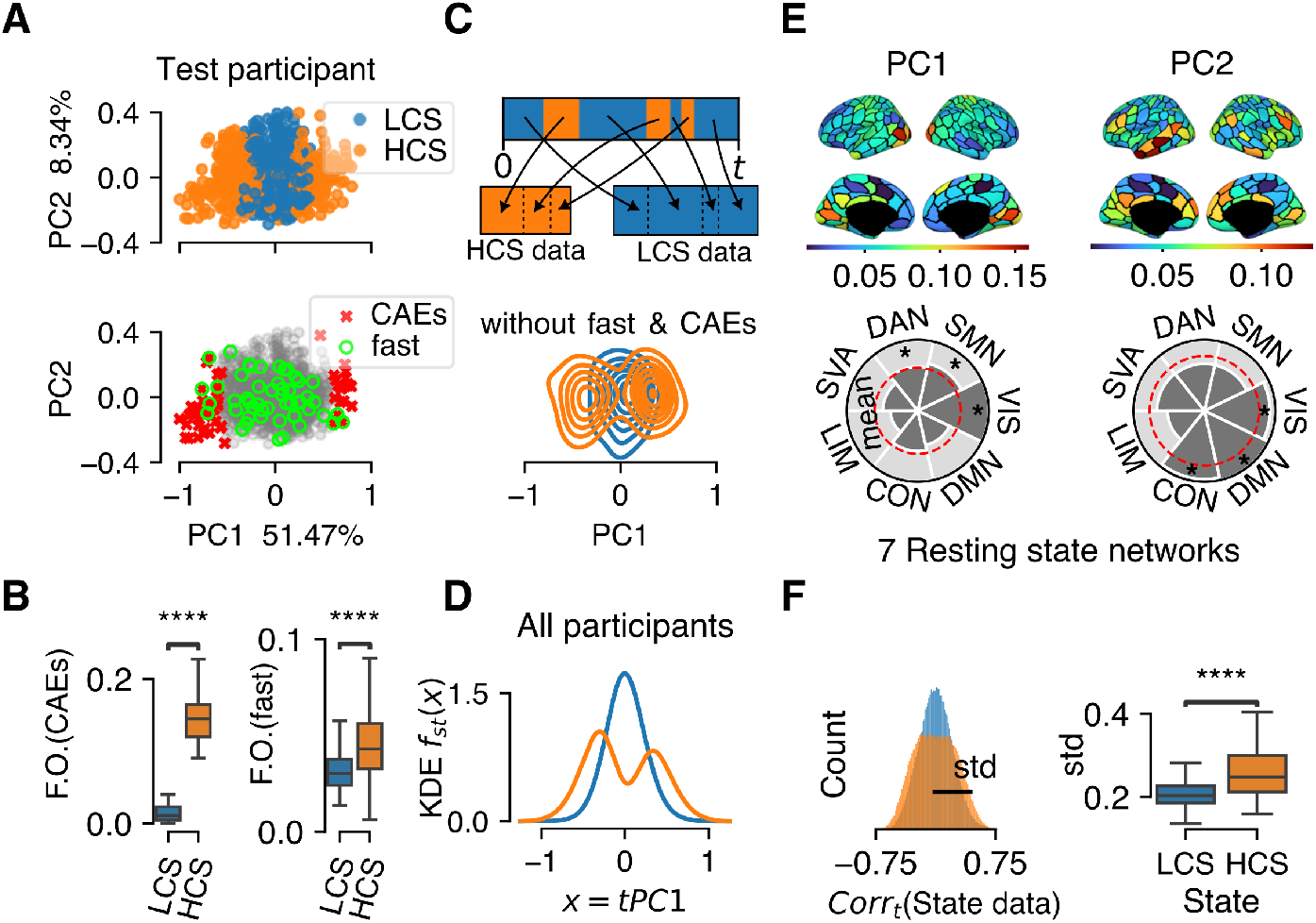
PCA reveals unimodal and bimodal stationary density functions for LCS and HCS, respectively, which translates into a richer variety of co-activation patterns during HCS. (**A**) The BOLD data were projected onto the first two PCs. The co-(de)activation events (CAEs) and fast time points were marked as transitional, non-stationary events. (**B**) The fast and CAEs fell strongly into the HCS. (**C**) HCS and LCS data were formed after concatenating their corresponding time points. After the removal of fast and CAEs, approximate stationary density functions (*f*_*st*_) were computed. (**D**) Fitting the probability densities using kernel density estimation (KDE) across the PC1 for all participants’ concatenated data, which revealed a unimodal and a bimodal distribution for LCS and HCS, respectively. (**E**) The spatial loadings of PC1 and PC2 were mapped to 7 resting-state networks. The VIS, SMN and DAN stood out with regard to PC1, whereas the DMN, CON and again VIS networks associated with PC2. (**F**) The standard deviation of the correlation of states data across time estimated the richness of co-activation patterns (Fig. S4). The higher variability of HCS along PC1 driven by the aforementioned networks conclusively equated with richer patterns of activity (*p* < 0.0001).

Aiming to get disassociated densities for the two states, we concatenated together the LCS and HCS epochs separately, forming the HCS and LCS stacks of BOLD data (Fig. 2C top). We removed the fast and CAEs and then estimated the stationary density functions for each state using kernel density estimation (KDE) (Methods D). For a test participant, the difference between the states came from their PC1 projection (Fig. S3A), which dominantly accounts for the largest amount of variance. This displayed a characteristic unimodality for LCS and bimodality for HCS, signifying shrinkage of activity between the two modes for the latter (Fig. 2C bottom). Focusing thereafter on the PC1, this picture was validated across all participants’ concatenated states data (Fig. 2D), demonstrating the existence of significantly different generative processes between the two states (KS test, *D* = 0.77, *p* < 10^−4^) (Fig. S3B). In addition, the regions’ spatial loadings (weights) of the PC1 (and PC2) of the overall activity were mapped to 7 RSNs (Thomas Yeo et al., 2011) (Fig. S6) using a mixed effects model (Bouzigues et al., 2024) (Fig. 2E). This suggested that functionally the difference is primarily driven by the peripheral networks —visual (VIS) and somato-motor (SMN) —and the dorsal attention network (DAN); and secondarily by the salient ventral attention (SVA), and the core networks —central executive control (CON), default mode network (DMN) and limbic (LIM). The PC2 loadings were stronger for the DMN, CON and VIS networks (Fig. 2E bottom). As a result, the PC1 axis delineated the familiar posterior-anterior gradient (Fig. 2E top-left) (Margulies et al., 2016); and thus, the manifold was situated primarily based on the linear combination of the peripheral networks and DAN. This explains the bimodality of HCS as a compressed form of co-(de)activations mainly of these resting-state networks (RSNs).

Finally, the richness of coactivation patterns was quantified for each state by the standard deviation of the correlation across time of the states data (Fig. 2F left), supported by a subsequent sensitivity analysis (Fig. S4A, B). Hence, the HCS marked significantly higher standard deviation than the LCS (*p* < 10^−4^) and thus entailed a richer variety of coactivation patterns despite its aforementioned short-lived nature.

### Bayesian hierarchical modeling identifies potential function parameters that characterize dynamics of the RSNs

The varying spatial contributions of the different RSNs to the PCs suggested differences in their underlying dynamics. To investigate this, we isolated the RSN-specific time series for each state and extracted their stationary densities, *f*_*st*_, from the PC1 scores (tPC1). These densities, referred to as *observables*, displayed unimodal and bimodal distributions of varying strength for LCS and HCS of the RSNs, respectively (Fig. S5, Fig. 3A, left). Under the working hypothesis that there are gradient dynamics that associate the modes of these densities to attractors (Methods E), the stationary solution of the one-dimensional Fokker-Planck equation describes the underlying energy landscapes of the states (Fig. 3B, top) (Jirsa and Sheheitli, 2022; Friston et al., 2023), expressed as generalized potentials *V* (*x*) (Haken, 1977). Borrowing on the concepts of *active inference* (Friston, 2013) to add another level of interpretation of the potentials’ geometries (Discussion), we notice that *V* (*x*) is the information-theoretic *surprise, I*(*x*), that extensively enters into the *free energy principle* (Fig. 3C, Methods I) (Friston et al., 2023). Such context imposes a complexity-accuracy trade-off between alternative potentials that can fit the observables. Model comparison (Fig. S7, Methods I) singled out the normal form of the quartic potential function (Fig. 3D, Methods E) and explained bistability as the sufficient complexity needed for the RSNs PC1 dynamics.

**Figure 3.**
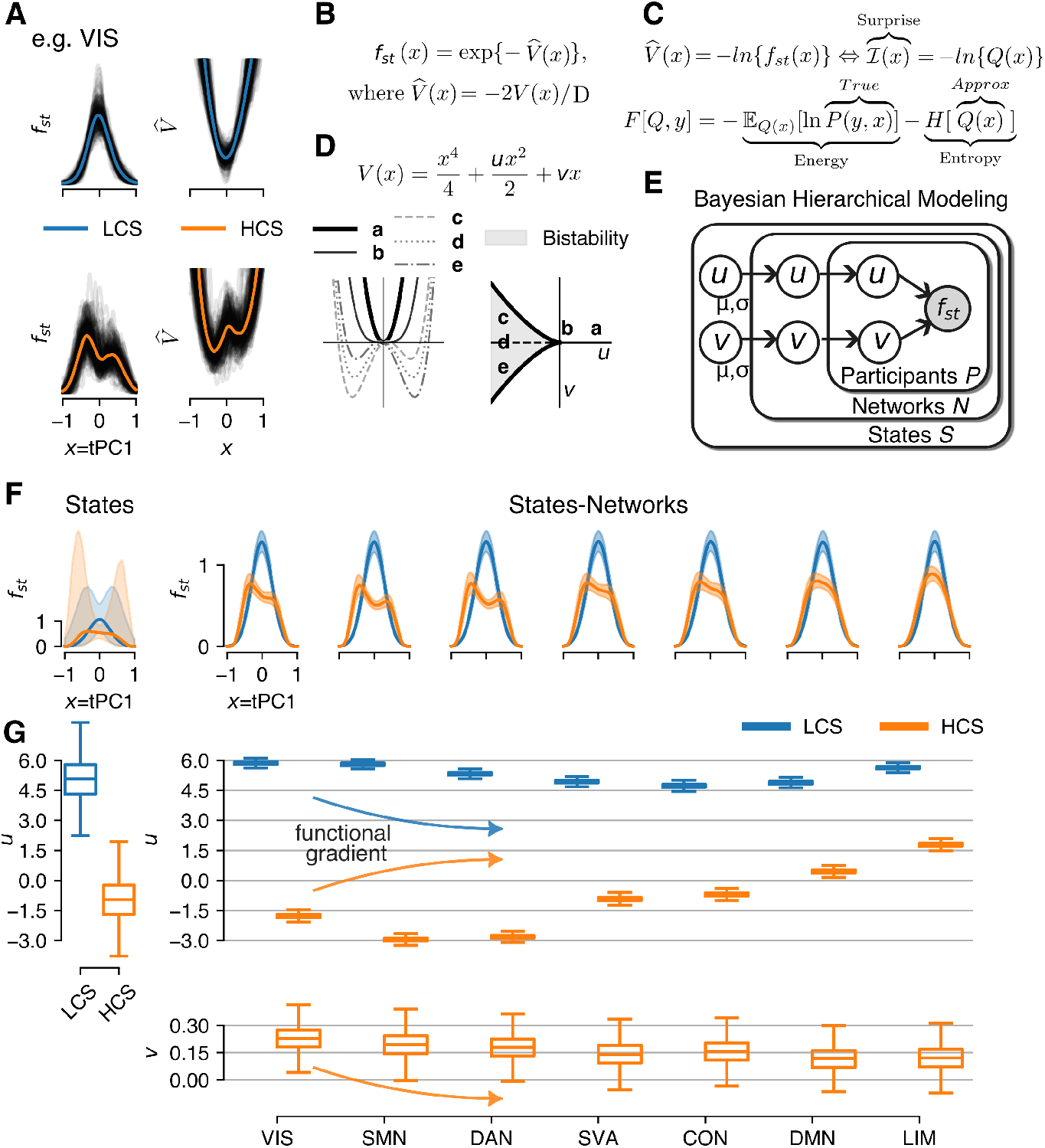
Bayesian hierarchical modeling fits quartic potential functions to explain the LCS (monostability) and HCS (bistability) stationary dynamics in the networks and states level. (**A**) Stationary density functions (observables) were extracted for each participant, network, and state, that reflect the underlying energy landscapes or potential functions 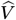. (**B**) This transformation stems from the solution of the stationary Fokker-Planck density function, where 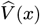 is the generalized potential function that absorbs both deterministic (*V* (*x*)) and stochastic (*D*) density forces. (**C**) The functional form of 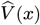 is equivalent to the information-theoretic *surprise, I* (*x*), which enters into the free-energy principle in the entropic term. (**D**) The normal form of the quartic potential function is sufficient to explain the geometry of the observables, with only two parameters, *u* and *v*. Here, *u* is the criticality parameter, which turns the system from monostable (*u >* 0) to bistable (*u* < 0), for *v* ~ 0; *v* is the bias parameter, which regulates the symmetry between the two wells, for *u* ≲ 0. The *u, v* combination creates a cusp bifurcation. (**E**) A Bayesian hierarchical modeling approach enabled inference of the parameters, *u*_*µ,σ*_, *v*_*µ,σ*_, at the *States-Networks-Participants* (same level as observables), *States-Networks*, and *States* levels, characterizing the latent dynamics. (**F**) Posterior parameter distributions were used to compute the posterior densities *f*_*st*_ of PC1 scores for each hierarchical level, shown in conformity with the observables (Fig. S5A). (**G**) Left: The geometrical difference of the states was now quantified by the *u* parameter which took large positive values for LCS (sharp monostability) and negative or positive and close to zero for HCS (bistability or flat monostability). Right-top: The parameter *u* noted a decrease in LCS that followed the functional gradient. This effect was mirrored and enhanced in HCS, as *u* was in the negative range for all RSNs except for DMN and LIM. Right-bottom: The bias parameter *v* also followed the functional gradient and was largest for the VIS network.

Bayesian hierarchical modeling was used to fit the observables, which allowed us to infer both individual- and group-level latent dynamical parameters across states and networks (Fig. 3F, G). Namely, the posterior distributions of two lower level parameters, *u* and *v*, were determined, that gave rise to the posterior densities *f*_*st*_ (Fig. 3F). At the *states* level, the LCS was sharply monostable (*u* ≫ 0), while the HCS exhibited a spectrum from flat monostability (*u* ~ 0) to bistability (*u* < 0) (Fig. 3G left), which is summarized as:

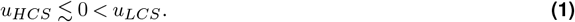

This clear differentiation between the states was explicated by the dynamics of the RSNs at the *states-networks* level (Fig. 3G right). LCS was again sharply monostable, with *u* ≫ 0 and a noticeable decrease that followed the primary functional gradient (Margulies et al., 2016). HCS mirrored and enhanced this effect, as confirmed by a significant correlation (*r*_*s*_ = − 0.79, *p* = 0.04) with the average T1w/T2w map, aggregated across networks (Fig. S8A-top). While *u*_*HCS*_ was mostly negative, it reached near-zero positive values in the DMN and rose well above zero in the LIM. Similarly, SVA and CON values were centered closer to zero. Additionally, *u* dominantly determined the characteristic relaxation timescale of a network close to its fixed point, for *v* ~ 0 (Fig. 3G bottom). For HCS, the pairwise timescales ratio between all networks was calculated (Methods F) and revealed that the DMN acquired the slowest timescale, which was larger than a factor of 10 between peripheral networks and DAN, and smaller for SVA and the other core networks (Fig. S10A). This result was only qualitatively replicated by the day-2 data analysis, as the DMN scored the slowest timescale but not larger than a factor of 10 (Fig. S10B). Together with dimensionality reduction (Fig. S1) and instabilities (Eq. (1)), the noticed timescales hierarchy provide empirical evidence for self-organization during the HCS, orchestrated by the DMN (Smallwood et al., 2021).

Secondarily, the bias parameter, *v*, also followed the functional gradient during HCS (Fig. 3G bottom, Fig. S9 bottom, Fig. S8A-bottom)and was the largest for the visual network. Noticeably, *v* was generally positive, which revealed the negative or low-activity solution as the favorable one between the two modes (co-deactivations), given bistability (Fig. 3B); or tilted the solution to the left, given flat monostability.

The third dataset was further used to test the biomarker potential of the inferred parameters. Power analysis of one-way balanced analysis of variance (ANOVA) indicated that for a large effect (*η*^2^ = 0.14) the number of participants required per group is 20.8; thus, the AD patients were not included in the analysis. In LCS, for both parameters there was no separation between CN and MCI, except for the SMN (*p* < 10^−2^) (Fig. S11A). In HCS, the *u* for the peripheral networks and the SVA significantly distinguished MCI from CN (*p* < 10^−2^) (Fig. S11B). Additionally, the effect sizes for *u*_*HCS*_ were all positive and larger than *u*_*LCS*_, except for SMN. The *v*_*HCS*_ also did not significantly separate MCI from CN for any network and noted negative effect sizes, but overall smaller than *v*_*LCS*_, except for VIS and DMN (Table S1). Last, the *u*_*HCS*_ for CN showed similar behavior across the RSNs (Fig. S11B-top, pink boxplots) as previously observed.

### The resting-state exhibits attractor manifolds that result from the deformation of the energy landscape alternating between the two states

SFMs and timescale hierarchies have been predicted from symmetry breaking in bistable neural masses (Perdikis et al., 2011; Woodman and Jirsa, 2013; Pillai and Jirsa, 2017). These properties are indirectly captured by their associated stationary densities, which change in response to the variation of a connectivity parameter, denoted by *G* (Fig. S12 and Methods G) (Jirsa and Sheheitli, 2022). We extracted the flow over the two-dimensional, PC1-PC2, manifold for a test participant (Methods J), demonstrating the attractor manifolds across the RSNs for both states (Fig. 4A). LCS exhibited a single attractor manifold (Fig. 4A, top), following *u*_*LCS*_ (Fig. 3G). In contrast, HCS was characterized by two attractor manifolds that progressively converged with respect to the functional gradient (Fig. 4A, bottom), predicted by the parameter *u*_*HCS*_ (Fig. 3G); similarly with the effect of *G* (Fig. S12). Figure 4A also indicated the existence of different decay rates associated with the flows across PC1 and PC2. We repeated the flow analysis for twenty participants and quantified that the dynamics along PC2 followed a slower, by a factor of 3, manifold compared to the fast convergence across PC1, for both states (Fig. S13B) (Methods K), with unrelated trend to the PC loadings (Fig. S13C). For reference, the network densities from all participants together were extracted, which supported a similar picture with Figure 4A (Fig. S13A).

**Figure 4.**
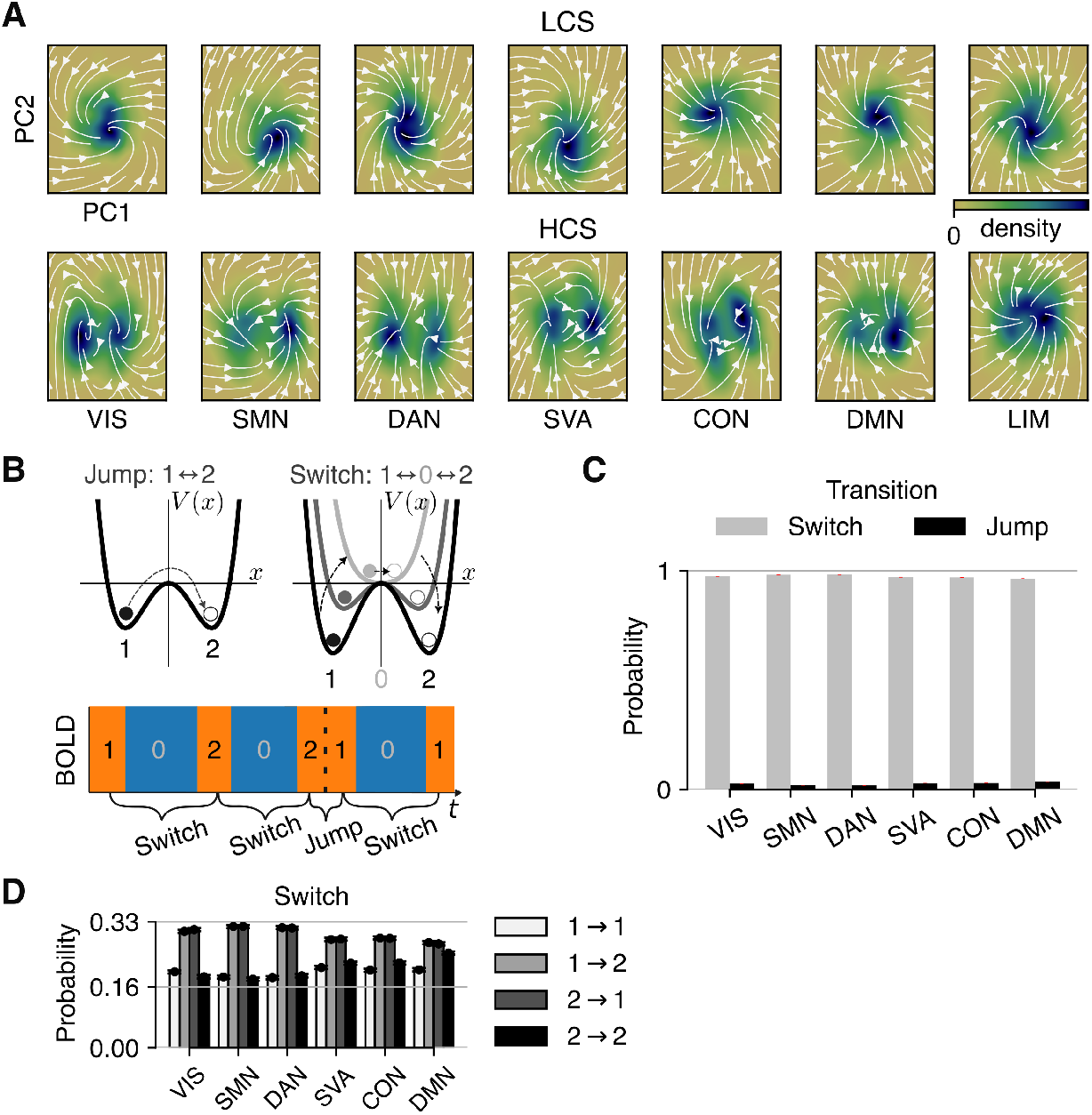
Dynamically changing attractor manifolds emerge following the alternation between LCS and HCS. (**A**) For an exemplary test participant, the flow was extracted, which illustrates the resting-state attractor manifolds for both LCS and HCS. The bistability in HCS predicts the generation of SFMs, which effectively are *jumps* between the two attractor manifolds. During LCS the dynamics are governed by a single attractor manifold (**B**) Top: illustration of the two possible mechanisms for transitions between modes: jump (direct transition) and switch (via deformation of the potential function). Bottom: schematic of how these mechanisms can be realized in the data, which led to classification of HCS datapoints into modes 1 and 2. (**C**) Switch is the dominant transition mechanism with a jump probability of only ~5%. (**D**) Transition probabilities between modes 1 and 2 are not symmetric, as state change is more preferable after a switch. This preference in switches weakens for the core networks.

So far, we have used time-independent stationary density functions to interrogate the state space dynamics of the resting-state, but how the system transitions between states of activity remains a question. Given that the system is at mode 1 (left mode) in the HCS, two options exist: either a *jump*, which effectively corresponds to an SFM and takes the system directly to mode 2; or a *switch*, which first deforms the potential function in order to bring the system to mode 0, or LCS, and then transition to mode 2 (Fig. 4B). Then, only for HCS, we classified every time point into its corresponding mode separately for each RSN (Fig. 4B-bottom, Methods B), excluding LIM that exhibited monostability. Given the limitations of the fMRI time resolution, we counted the number of jumps and switches to estimate the frequentist probability of a switch against a jump which revealed that jumps happened only at 5% of all the transitions that took place (Fig. 4C). The transitions did not exhibit symmetrical behavior as there is a preference for a mode change after either a switch or a jump (Fig. 4D), meaning that mostly co-activations follow co-deactivations and vice versa. For the switches, this preference fades following the functional gradient, brought by *u* and *v* that shape the geometry of the dynamics (Fig. 3G).

A further qualitative sensitivity analysis used the above transition probabilities as transition matrices that tuned the dynamics of a tri-stable periodic model very close to a threshold (bifurcation) where only one solution would survive (S14A, B) (see Methods L). This suggested that the empirical system may operate near a bifurcation point (metastability), where small changes can turn it from multistable to monostable.

### CAEs provide a mechanism for jumps across PC1 axis or traversing across the slow manifold along the PC2 axis

Previous works have indicated that CAEs can be driven by neuronal cascades and, mechanistically, can facilitate nodal jumps between two stable states of whole-brain bistable neural mass models (Rabuffo et al., 2021; Fousek et al., 2024); thus, we focused on their influence on the previously identified jumps. Figure 5A displays an example of a HCS trajectory that exhibits such a jump accompanied by CAEs. This exact trajectory is further projected on the PCs space for clarity (Fig. 5D). Interestingly, almost 50% of all jumps were driven by CAEs (Fig. 5B). The difference with jumps that occurred without CAEs lied on the distance covered across the PC1 (PC1-translation), which was significantly lower than with CAEs (Fig. 5F left). This meant that such trajectories moved around zero, implying that the potential barrier might have been lower, so that CAEs were unnecessary to escort the trajectory to the other state (Fig. S16). However, jumps happened very rarely and explained only a small fraction of all CAEs. Figure 5C demonstrated HCS activity that involved CAEs but no jump. Such trajectories are, thus, mode-locked. The projection on the low-dimensional PCs manifold demonstrated a long excursion along PC2 (Fig. 5E). However, the dynamics along PC2 involve a slow manifold as previously shown by the flow (Fig. 4C, Fig. S13B). This difference was quantified between the rest of the trajectories involving CAEs and without it (Fig. 5F right), and likewise explained that the CAEs promote longer exploratory trajectories defying the canonical flow on the resting-state manifold.

**Figure 5.**
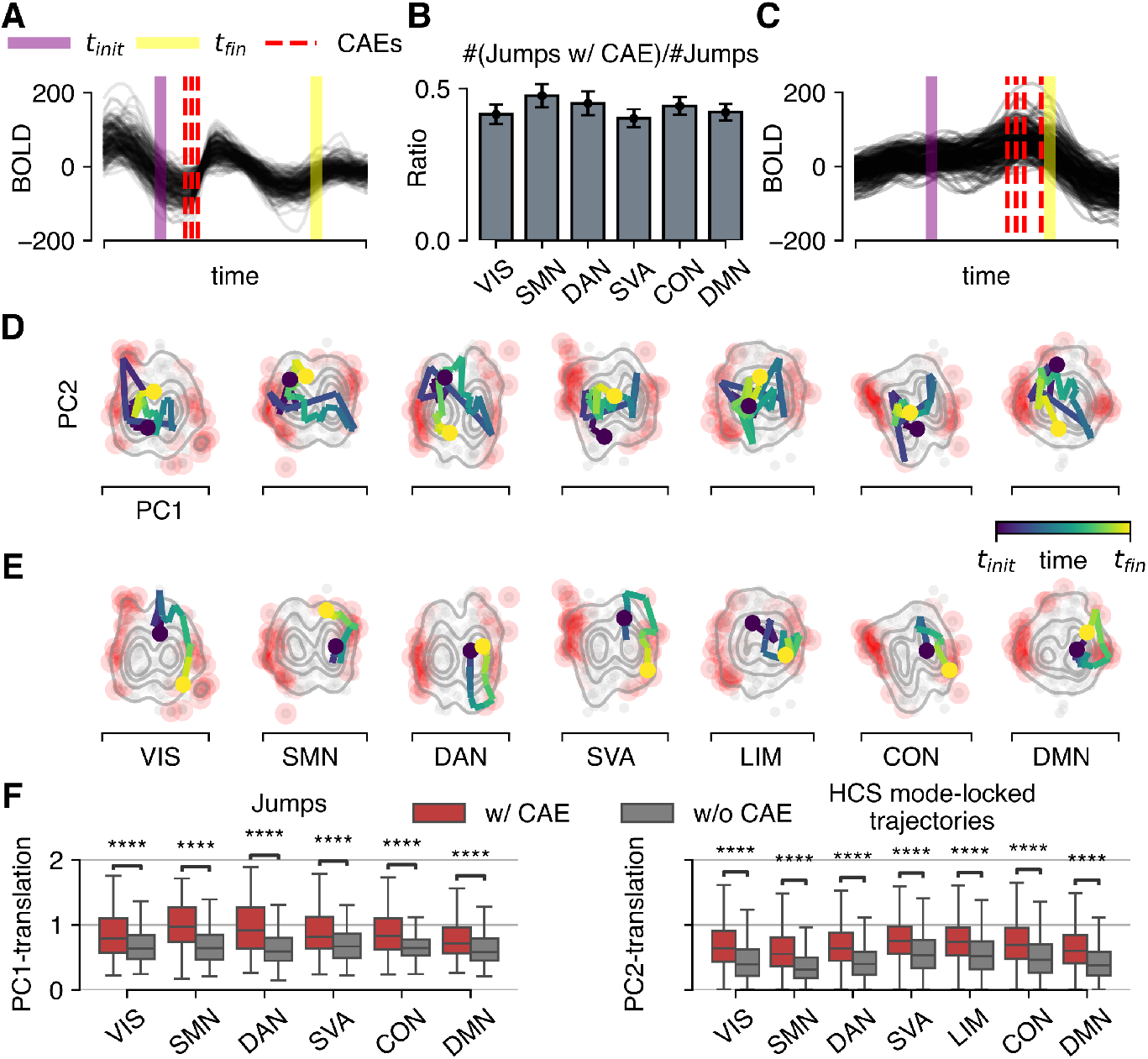
In HCS, CAEs are stepping stones for cross-mode jumps across PC1 or traversing along the slow manifold in PC2. (**A**) HCS BOLD activity that exhibits cross-mode jump starts at *t*_*init*_ and finishes at *t*_*fin*_. The jump is accompanied by CAEs. (**B**) Almost 50% of all the jumps took place together with CAEs. (**C**) Similar to **(A)**, but while CAEs happen, no jump occurs. (**D**) The low-dimensional projection of the *t*_*init*_ to *t*_*fin*_ activity in (**A**) is visualized and demonstrates the cross-mode jump. (**E**) Same as (**D**), but taken from the activity in (**C**), which demonstrates an extended trajectory along the slow manifold of PC2. (**F**) The CAEs promote longer exploratory trajectories whenever a jump (left), or else mode-locked activity across PC2 happens (right).

## Discussion

Here, we have demonstrated how the labile brain (Friston, 2000) is embodied in the slow dynamics of the BOLD signal through a dynamically changing resting-state attractor manifold: from unimodal, hosting incoherent activity (the LCS), to bimodal, with two attractors, to facilitate high coherence dynamics (the HCS) (Fig. 2, Fig. 4). This transition entailed a reduction in effective dimensionality (Fig. S1) and a reorganization of the RSNs dynamics (Fig. 3G, Eq. (1)) consistent with the functional gradient (Margulies et al., 2016), and temporal hierarchies that revealed slow timescales for SVA and core networks. Together, these changes provide empirical evidence of self-organization during HCS, substantially orchestrated by the DMN, which noted the slowest timescale (Fig. S10). Although HCS was short-lived compared to the dominant LCS (Fig. 1), it was marked by more diverse co-activation patterns and hosted the strongest non-stationarities (fast and CAEs) of the BOLD signal (Fig. 2). Notably, CAEs facilitated dynamically improbable flows in the HCS, including cross-mode jumps across PC1 and extensive exploratory trajectories along the slower manifold in PC2 (Fig. 5).

In contrast to numerous studies focusing on fine-grained coactivation patterns to explain resting-state organization (Vidaurre et al., 2017; Peng et al., 2023), our coarse-grained coherence analysis implicitly assembled patterns corresponding to similar subspaces within a low-dimensional manifold. The division of spontaneous BOLD signal into LCS and HCS has been noted previously (Zalesky et al., 2014; Kong et al., 2021); however, our approach employed a temporally fine-grained method to extract these states based on instantaneous relative phases (Fig. 1A, B). While prior studies have used similar phase-based methods, they adopted different clustering approaches at the phases eigenspace that did not characterize the resting-state manifold, yet they successfully identified distinct brain states for various applications (Cabral et al., 2017; Kringelbach et al., 2020).

In this work, projection onto PC1 alone sufficed to distinguish the manifold organization of LCS and HCS. The intriguing bimodality of the HCS was spatially associated with activity in peripheral networks (VIS and SMN) and the DAN (Fig. 2E), rather than the SVA and core networks —typically considered the primary drivers of resting activity (Christoff et al., 2016). However, SVA, CON and DMN were explicitly found to be the closest to zero (Fig. 3G), or *instabilities*, which renders them as the “unstable modes”, contrasting the other networks that can be referred to as “stable modes” (Haken, 1977). Technically, the requisite time-scale separation appears to be manifested only by the DMN (Fig. S10), which can therefore be considered to orchestrate HCS activity by imposing its slow dynamics on the periphery and the DAN, thereby materializing the self-organization of brain activity across the cortical hierarchy (Smallwood et al., 2021). Moreover, since the CON is functionally associated with deliberate control over attentional shifts, and the SVA with rapid, spontaneous shifts of attention, these networks are appropriately found close to instabilities. In contrast, the DAN is considered to stabilize attention (Christoff et al., 2016), and sensory or motor areas are associated with stimulus-/task-locked activity even in the absence of overt activity during rest (Raichle, 2011).

Methodologically, the use of the stationary probability densities derived from the solution of the Fokker-Planck equation has been instrumental in our exploration of the underlying dynamics. In random dynamical systems, these densities imply the existence of pullback attractors of the states (Crauel and Flandoli, 1994; Friston et al., 2023), which we identified as the attractor manifolds for LCS and HCS (Fig. 4). In analogy with Jirsa and Sheheitli (2022), the HCS parameters systematically altered the manifolds’ geometry along the functional gradient, exhibiting intra-network connectivity properties (Fig. S8). Incorporating PC2, a timescales difference was also revealed: fast convergence across PC1 and slower dynamics across PC2 (Fig. 4C, Fig. S13B). This situated the fixed points within slow manifolds, which topologically can be equivalent to approximate continuous attractors (Ságodi et al., 2024). These are understood as pattern-forming continuous-attractors, which can enable networks to integrate and encode information over long timescales (1-100 s) (Khona and Fiete, 2022). Given the presence of instabilities (symmetry breaking) and noise, these densities may reflect SFMs that govern trajectories between different modes (Fig. S12) (Jirsa and Sheheitli, 2022). Here, SFMs manifested as rare jumps between HCS modes. More dominantly though, transitions occurred via potential function deformation (switch), which is known under the Landauer’s principle (Landauer, 1961) to require entropy production. The global coordination seen in HCS indicates a low-entropy configuration that dissipates into the incoherence and higher entropy of LCS. This switching mechanism enables information storage (consolidation) and preparation for subsequent transitions (Clawson et al., 2019), whereas diffusion-driven jumps are metabolically expensive, less reliable, and associated with computational memory loss (Haken, 1977; Bennett, 1982). Nonetheless, the presence of jumps provides evidence for the co-existence of two modes during HCS, consistent with the stationary densities. The participation of CAEs —the strongest BOLD co-activations and thus most metabolically costly —during such jumps (Fig. 5) reinforces the notion that switching via potential deformation is not only computationally advantageous, but also metabolically cheaper.

An additional layer of interpretation arises from the equivalence between the stationary Fokker–Planck equation and surprise in information theory. Stationary density functions can then be viewed as marginalized priors shaped by active inference (Fig. 3C, Methods I Eq. (10)) (Friston, 2013; Dimakou et al., 2025). In this framing, the networks’ dichotomy observed above can be represented by sensory and internal states, respectively (Parr et al., 2022). The parameter *u* determined the geometry of internal state distributions, linking them to the type of information sampled from sensory states under the principle of maximum entropy (Jaynes, 1957, 2007) (second term in Eq. (11), Fig. 3C). During HCS, sensory states acquired a bimodal distribution, while internal states approached a uniform distribution (Fig. 3F), which maximizes entropy for discrete random variables (Bishop, 2006). This suggests that information encoded by internal states during HCS is discrete. Conversely, during LCS, both sensory (higher precision) and internal states followed unimodal, Gaussian distributions (Fig. 3F), which maximize entropy for continuous variables (with constrained first two moments) (Bishop, 2006), suggesting that the information is of *continuous* nature. This explanation can link to the findings from Schroeder and Lakatos (2009), which distinguished either a rhythmic or a continuous (or vigilance) mode during sensory selection. The former requires a temporal structure that is entrained by low-frequency oscillations (associated with BOLD infra-slow frequencies (Monto et al., 2008)). The latter takes place in the absence of any rhythm that the system can entrain to, and gamma oscillations are dominant. At the same time, the nonzero bias parameter, *v*, speaks for the *Energy* term (Fig. 3C, Eq. (11)), which assigns favorable prior states, or implies that the generative internal model anticipates the prior sensations to be most likely satisfied (*optimism bias*) (Friston, 2013). This may explain why the visual network showed the strongest bias during HCS (Fig. 3G bottom), given participants’ eyes-open and relaxed fixation instructions (Methods A).

Integrating our results with previous literature, we can summarize the following arguments pointing toward a specific interpretation of LCS and HCS: **1**. HCS is characterized by short-lived trajectories of highly correlated, synchronized, and thus coordinated signal, reminiscent of phases of *entrainement* (Lakatos et al., 2019); **2**. even though resting-state is dominated by the activity of DMN and other core networks which drive stimulus independent processes like mind-wandering or daydreaming (spontaneous thought) (Christoff et al., 2016), the SMN appears more mechanistically responsible for state transitions (or FCD switches) (Kong et al., 2021); **3**. a significant variation in RSN dynamics occurs from LCS to HCS, as peripheral networks transition from the overall least variable to most (Fig. 3G), highlighting a transition of the dynamics from unimodal to bimodal and therefore suggesting a potential shift in the system’s functional state; **4**. however, since jumps were rare, the activity during a single HCS trajectory evolved within a specific mode for all networks (e.g. Fig. 5E), implying lower variability than LCS trajectories evolving under the monostable distribution (Fig. S16). This aligns with the well-established observation that task/stimulus-driven activity exhibits lower signal variability than pre-stimulus periods across the cortex (He, 2013; Ponce-Alvarez et al., 2015; Wu et al., 2024). Taken together, these observations suggest that HCS resembles task-based activity (whether internally generated or driven by external interactions, i.e. active inference), whereas LCS reflects spontaneous thought (passive inference). This considers that the resting-state paradigm inherently involves an environment with external stimuli (e.g. scanner sounds, vibrations) that can engage sensory networks, producing fleeting shifts from task-free to task-based modes. Even without overt action, perception resolves prediction errors from mismatches between internal and external states (Parr et al., 2022).

Although LCS is the dominant dynamic at rest, a greater amount of time spent in HCS was associated with higher brain fluidity (Fig. 1D). Dynamically, fluidity is linked to the richness in the state space, which reflects the complexity of the functional repertoire (Breyton et al., 2023). In electroencephalographic recordings, whole-brain fluidity has emerged as a feature characterizing AD patients (Mazzara et al., 2025). However, for BOLD recordings in the third dataset, fluidity alone was insufficient to distinguish MCI from CN (Fig. S17). Similarly, the stereotypical, single-fixed point dynamics of the LCS across the RSNs, described mainly by the parameter *u*, did not show cohesive sensitivity to the alternated MCI brain hemodynamics (Owens et al., 2024), as only SMN displayed significantly different *u*. This finding underscores that the incoherent nature of resting-state fMRI data challenges the identification of robust features for distinguishing healthy controls from clinical populations. It was only during the HCS —which arguably reflects a form of task-based activity —that the richer, bimodal dynamics showed sensitivity to the MCI dynamics, displaying a robust positive drift for *u*, significant for the peripheral and attention networks and overall larger effect sizes (Fig. S11, Table S1). However, the short-lived nature of HCS necessitates a high rate of image acquisition and long scanning sessions to ensure sufficient statistical power that effectively displays the dynamics of the stationary densities. The datasets used here provided the necessary specifications for this temporally fine analysis, yielding similar fluidity correlations (Fig. 1D and Fig. S2C, D) and scaling of the *u* and *v* parameters (Fig. 3G, Fig. S9 and Fig. S11).

Fundamentally, the inherent slowness of the fMRI signal limits our dynamical interpretations, as it only allows for observations of putative fixed points. Consequently, the coexistent bimodality during HCS and the nature of state transitions are also constrained by the true timescale of the underlying dynamics: jumps must occur faster than switches to be distinguishable. Analysis with faster time-resolution electrophysiological recordings could provide more accurate insights into the non-linear dynamics of these states, and is necessary to compare with our interpretations. Nonetheless, our focus on stationary dynamics helped mitigate this issue, offering a baseline characterization of the resting-state dynamics and manifold as a function of signal coherence. Building on this, future work should investigate the relationship between HCS and arousal levels during rest, which have been shown to modulate BOLD fluctuations (Raut et al., 2021). Active inference modeling could be employed to link arousal, coherence, and dynamical control parameters hierarchically (Parr et al., 2022), capturing the evolution of resting-state activity in a holistic manner. Thus, we hope this work lays the groundwork for new strategies in resting-state whole-brain modeling that incorporate these mechanistic insights. Finally, the link to task-based activity calls for a direct comparative analysis with task-fMRI data to further explore the dynamical-active inference interpretation we attributed to the control parameters, *u* and *v*. Such analysis will be crucial for characterizing their nature and evaluating their potential value as biomarkers.

## Methods

### A. Data

Data used in preparation of this work was initially obtained from the Human Connectome Project (HCP) dataset (Van Essen et al., 2012), whose details are published elsewhere (Smith et al., 2013). A subset of 200 young healthy participants (96 males), resting-state BOLD data from the left-right encoding session were used, consisting of 1200 time points, sampled at 720 ms, parcellated to the Schaeffer 200-parcel 17-network brain atlas (Schaefer et al., 2018). The data is available in the EBRAINS repository https://doi.org/10.25493/F9DP-WCQ (Domhof et al., 2021). Here, we aggregated the areas into the most prominent 7 resting-state networks to perform the networks analysis, and disregarded the temporal-parietal junction network stemmed from the original 17-networks atlas.

Data used in the preparation of this article were obtained from the Alzheimer’s Disease Neuroimaging Initiative (ADNI) database (adni.loni.usc.edu). The ADNI was launched in 2003 as a public-private partnership, led by Principal Investigator Michael W. Weiner, MD. The primary goal of ADNI has been to test whether serial magnetic resonance imaging (MRI), positron emission tomography (PET), other biological markers, and clinical and neuropsychological assessment can be combined to measure the progression of mild cognitive impairment (MCI) and early Alzheimer’s disease (AD). A high-resolution (TR=0.6) subset (48 participants) of the ADNI dataset made up the third dataset. The minimal preprocessing pipeline from HCP (Glasser et al., 2013) was applied to preprocess the functional imaging data. For the preprocessing we used the fMRIVolume pipeline, which included correction for head motion, susceptibility-induced distortions, and bias field inhomogeneities, followed by alignment to the corresponding T1-weighted structural image. The subsequent fMRISurface step involved resampling the data onto each subject’s grayordinate space—comprising both cortical surfaces and subcortical volumes—and applying spatial smoothing using a Gaussian kernel (FWHM = 2 mm). To denoise the data, we trained FSL’s FIX classifier using a manually labeled dataset of neural and artefactual independent components from 23 subjects (Salimi-Khorshidi et al., 2014). This trained classifier was then applied to all subjects to remove artefactual components from the fMRI data. Finally, the cleaned time series were parcellated using the Schaeffer 100-parcel, 7-network brain atlas to extract region-wise time series data.

### B. States identification, fluidity and temporal metrics

LCS and HCS data points were identified using a time instantaneous approach, adapted by Cabral et al. (2017). The signal was first bandpass filtered from 0.01 to 0.18Hz (Gonzalez-Castillo et al., 2015), and a Hilbert transformation yielded the analytical signal, namely complex vectors. The pairwise cosine differences of the vectors’ angles was calculated for each time point to generate instantaneous phase coherence matrices of shape NxNxT, where N is the number of regions of interest (ROIs) or nodes, and T the number of samples. The collection of the vectorized upper triangle of the NxN matrices provided us with a two-dimensional matrix of shape N(N-1)/2xT. This matrix was used to calculate, first, the mean coherence, by considering its absolute values and averaging across the first dimension (Fig. 1B top); second, fluidity, by calculating the correlation across time and then the variance of the upper triangle of that correlation matrix for each participant (Fig. 1D). The one-dimensional mean coherence array of shape T was used as input to a Hidden Markov model (HMM) (Vidaurre et al., 2017) with two hidden states to separate between global LCS and HCS. Importantly, the HMM accounts for the time-dependencies of the data which is an important consideration for the formation of states. Fractional occupancy of the states was calculated by dividing the number of all occurrences by the total number of time points of the signal. Mean dwell times were the average duration of each state occurrence before transitioning to the other state. Specifically, the high-cut of the bandpass filter was selected such that the correlation between fractional occupancy and fluidity across participants maximized (Fig. S2A). After isolating the HCS data (Fig. 2C-top), we performed a second HMM analysis to classify the HCS datapoints into modes 1 and 2 (Fig. 4B).

### C. Non-stationarities identification

The CAEs mark extremely high co-(de)activation across all areas that resulted in activity taking place at the boundaries of the manifold, denoting the strongest transients (Fig. 2A bottom). They are identified from the edge time-series defined as *E*_*nm*_(*t*) = *z*_*n*_(*t*)*z*_*m*_(*t*), for *n, m* = 1 … *N*, where 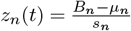 is the z-scored BOLD time-series of a node n. The CAEs are defined as time points in the edge time-series *E*_*nm*_(*t*) during which the root sum squared 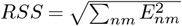 crosses a given threshold, here chosen as 95th percentile. Mechanistically they are considered to bring about transitions (Rabuffo et al., 2021; Fousek et al., 2024), as it was also illustrated in Figure 5. Second, we identified the fast time-points of activity as the ones with the highest speed, quantified by the mean Euclidean distance covered between two consecutive time-points in the PCs space (Fig. 2A bottom). Since both of these non-stationary events are presumably of similar nature, we retained as many PCs, i.e. 4, as necessary to maximize the overlap between these two classes of non-stationarities. The fast time points were identified by setting a threshold at three times the median absolute deviation of the speed of the vectors in that PCs space.

### D. Empirical stationary density functions

The spatial analysis of the states was performed on the unfiltered data. To approximate stationary probability distribution functions we excluded from the signal the non-stationarities and concatenated the segments corresponding to each state. We applied PCA first across all nodes and then on the individual seven RSNs (Thomas Yeo et al., 2011). Kernel density estimation (KDE) on the PC1 scores (tPC1) approximated the empirical probability density functions. We used a kernel bandwidth of 0.1 for the global density functions, which balanced the local and global variation of activity; a smaller bandwidth of 0.075 for the networks activity was used to promote localized variation, which intended to highlight the differences between networks (Fig. S5) at the cost of smoothness. Additionally, lower and higher bandwidths were used (0.05, 0.066, 0.09), which did not qualitatively alter the results.

### E. Potential functions

The temporal PCs scores (*x* =tPC1) were treated as the state variables of the system (Fousek et al., 2024), which admit deterministic and stochastic forces. This is expressed by a Langevin equation, which, in the one-dimensional form, reads as:

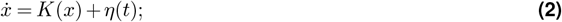

where *K*(*x*) and *η*(*t*) correspond to the deterministic and stochastic component, respectively. The Fokker-Planck equation captures the time evolution of the probability density dynamics of *x*, i.e., *f* (*x*), incorporating these components. The so-called potential condition (Stratonovich, 1967) for the one-dimensional case is the “natural boundary condition” (Haken, 1977), which requires that *f* vanishes for *x* → ±∞, and:

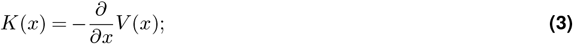

where the deterministic force is expressed via the negative gradient of the underlying potential function. The stationary solution of the Fokker-Planck equation renders the statistical properties of the states (Jirsa and Sheheitli, 2022) and reads as:

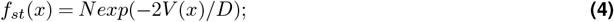

where *N* is the normalization constant and *D* is the diffusion constant that stems from *η*(*t*). Further on, Eq. (4) can be rewritten as:

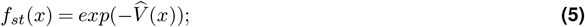

where 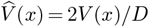, as 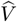 absorbs *D* (Fig. 3B top) (Haken, 1977). Hence, we can transform the *f*_*st*_ observables into generalized potential functions, 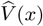 (Fig. 3A right), which allowed us to think of possible functional forms that can explain our observables. We selected the normal form of the quartic potential function (*Thom’s unfoldings* (Zeeman, 1979)), which gets rid of superfluous terms:

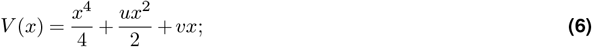

Eq. (6) is comprised of only two parameters, i.e. *u* and *v. u* principally determined the stability of the state space, while *v* controlled for the asymmetry of the two wells, given bistability (*u* < 0) (Fig. 3B-bottom-left). Therefore, *u* is termed as the criticality parameter and *v* the bias parameter. The combination of the *u, v* control parameters makes up a cusp bifurcation (co-dimension 2) in the state space which determines whether the system is monostable or bistable (Fig. 3B-bottom-right).

### F. Timescales hierarchies and self-organization

Combining Eq. (3) and Eq. (6), the corresponding equation of motion is:

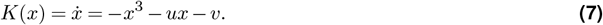

For *v* ~ 0 (Fig. 3-right-bottom) the stationary solution of Eq. (7), i.e. *K*(*x*) = 0, is determined by the parameter *u*, as the stable fixed points are at *x*_∗_ = 0, for *u >* 0, and 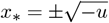, while *x*_∗_ = 0 now becomes an unstable solution. In both cases, the characteristic timescale of the relaxation around the fixed point is determined by the value 1*/*|*K*^*′*^(*x*_∗_)| (Strogatz, 2024). This is:

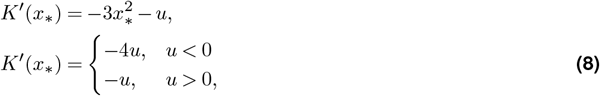

where we substituted in Eq. (8) the solutions mentioned above. Based on these cases the timescale of each network in the HCS was estimated from its mean *u* (shown graphically in Figure 3-right), namely *τ*_*RSN*_ = 1*/* | *K*^*′*^(*x*_∗_,⟨*u*_*RSN*_⟩) |. Then the tau ratios between networks can be computed (Fig. S10). Typically, if *τ*_*ratio*_ ≫ 1, then by an adiabatic approximation, self-organization predicts that the slow subsystem “enslaves” the dynamics of the fast subsystem imposing its slow dynamics. This dynamic takes place only if the slow modes operate close to instabilities (also known as unstable modes) (Haken, 1977), which happens in Eq. (7) close to the bifurcation point, *u* ~ 0.

### G. Symmetry breaking in bistable networks

Incorporating another dimension into the dynamics for each potentially bistable network (that would be in practice the temporal PC2 scores) makes up a coupled system of differential equations. A two-dimensional generalization of Eq. (7), together with the noise term *η*(*t*) from Eq. (2), is (Haken, 1977):

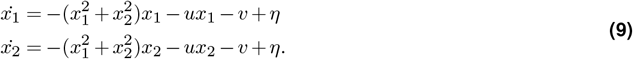

For *u >* 0 we end up again to a single fixed point. To examine the bistable case, we let *u* < 0 and for simplicity (toy example) we set it to *u* = −1. Moreover, to control the coupling *x*_1_, *x*_2_, which appears as 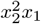 (and 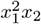,respectively) we introduce a complementary coupling term modulated by a factor *G* (Jirsa and Sheheitli, 2022); and Eq. (9) becomes

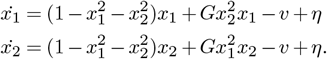

The networks dynamics are now effectively regulated by *G*, ranging from 0 to 1. We simulated this system for the symmetric bistability, i.e. *v* = 0 (Fig. S12). For *G* = 1, the dimensions become uncoupled as the coupling terms cancel each other out and the nullclines form infinite straight lines. For strong enough noise, the trajectory can escape from one attractor and very fast transition to another one, thus forming four attractors (two bistable nodes) in the space. To the other extreme, for *G* = 0 the topology becomes equivalent to all-to-all coupling and the nullclines converge to a continuous ring attractor manifold. The trajectory then lives within this manifold and its movement therein is fully stochastic. For intermediate values, e.g. *G* = 0.5, the nullclines are elliptic and trajectories exhibit structured flows on the manifold (SFMs) determined both by the deterministic and stochastic component inducing timescales separation between the flow and the collapse onto it (Jirsa and Sheheitli, 2022).

### H. Bayesian hierarchical modeling

The *f*_*st*_ observables from the tPC1s were normalized for each network separately prior to the Bayesian regression. A Bayesian Hierarchical Model (BHM) was employed to infer the latent dynamics of the *states* or root nodes of the graph, and the *states-networks* nodes given the observables *f*_*st*_ at the *states-networks-participants* level or root nodes (Fig. 3C). The BHM architecture required only to assign the mean and standard deviation (*µ, σ*) of the parameters, *u* and *v*, at the states level. It utilized the chain rule to propagate in a feed-forward manner the prior distributions of the parameters, while the geometry of the observables back-propagated from the leaf to the root nodes. As a result of the observables pooling, the non-leaves nodes’ posterior distributions did not suffer from the localized kernel bandwidth that we selected for the participants’ networks density functions. The inference was performed using adaptive Hamiltonian Monte-Carlo, No-U-Turn Sampling (NUTS) algorithm, implemented via the numpyro package (Bingham et al., 2019; Phan et al., 2019).

### I. Model selection

The generalized potential function from the solution of the stationary Fokker-Planck equation has the same functional form as the surprise from information theory:

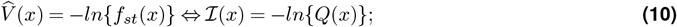

where ℐ (*x*) is the surprise, and the stationary density function correspondingly translates into the *approximate posterior* or *internal model* which compares to the *true posterior, P* (*x* | *y*), when interacting with (or sampling information from) the environment. From Bayes’ rule, *P* (*x* | *y*) = *P* (*y, x*)*/P* (*y*); and entropy is the long-term average of surprise, *H*[*Q*(*x*)] = 𝔼_*Q*(*x*)_[*I*(*x*)], which appears directly in the variational free energy equation:

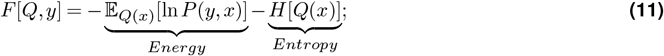

where *F* [*Q, y*] is intended to minimize with respect to the trade-off between the *Energy* and the *Entropy* terms. Namely, the selection of the potential function *V* (*x*), within the framework of active inference, comes naturally as a quest for an internal model that satisfies the balance between *entropy maximization*, i.e. Jaynes’ principle in complete absence of prior knowledge (Jaynes, 2007)), and *energy minimization*; as *Q*(*x*) intends to assign higher weights to states *x*, where the true model *P* (*y, x*) is large (inversely proportional to energy). Since Eq. (11) can be rewritten as:

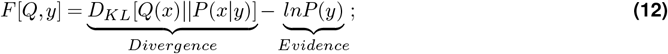

the quest to find the best *Q*(*x*), becomes complementary to maximizing the model log evidence. Approximations to negative model log evidence that can be used are the Bayesian information criterion (BIC) and Akaike information criterion (AIC) (Parr et al., 2022), which are functions of the estimated maximum likelihood (Akaike et al., 1973; Schwarz, 1978). Simpler (*P*_2_(*x*)) or more complex (*P*_6_(*x*), *P*_8_(*x*)) models can be candidates for fitting the observables. All polynomials used, followed their normal forms (Zeeman, 1979; Haken, 1977), namely:

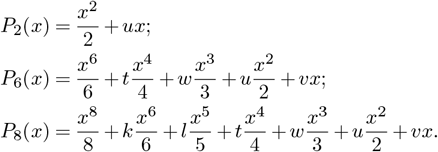

These models, together with *P*_4_(*x*) or Eq. (6), were compared based on the aforementioned information criteria.

### J. Flow extraction

The flow on the manifold for the states data (Fig. 2C top) is computed by fitting a recurrent switching linear dynamical system (rsLDS) (Nair et al., 2023) on the low-dimensional space as expressed by the first two PCs. The exemplary test participant exhibited balanced fractional occupancy between LCS and HCS which was important for the validity of the fitting.

Initially, for a test participant, rsLDS models (Linderman et al., 2017) were fit to the RSNs’ BOLD states data (Fig. 2C-top) separately for each network. The rsLDS model is a generative state-space model that uses piecewise linear models to fit the dynamics of complex nonlinear time series. Three types of variables describe these interacting dynamics: 1. a set of discrete states *z*_*t*_ ∈ {1,…, *K* }, which represent hidden modes or distinct regimes the system can occupy; 2. a set of continuous latent variables *x*_*t*_ ∈ ℝ ^*D*^, which capture the low-dimensional structure of the underlying process; 3. the observed activity *y*_*t*_ ∈ ℝ ^*N*^, representing the BOLD recordings from *N* nodes. We used two-dimensional models (*D* = 2) in consistency with (Vinograd et al., 2024), while we had a clear hypothesis for The bistability in peripheral networks in HCS suggested *K* = 2, and for the core networks we performed model evaluation via the evidence lower bound computation for *K* = 2, 3, 4, as explained in (Nair et al., 2023). For the trivial LCS monostable dynamics, we still used the same rsLDS model for consistency and validation purposes, with *K* = 2.

In particular, at each time step *t*, the model determined the next discrete state *z*_*t*+1_. This transition is not random; it’s dynamically influenced by the current continuous latent state *x*_*t*_. This relationship is formally expressed through a softmax function applied to a linear transformation:

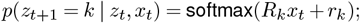

where, *R*_*k*_ ∈ ℝ^*K×K*^ and *r*_*k*_ ∈ ℝ^*K*^ are parameters that govern the probability of transitioning into state *k*, based on the current latent activity *x*_*t*_. Once a discrete state *z*_*t*_ is active, it defines a unique linear dynamical system that dictates how the continuous latent variables *x*_*t*_ evolve:

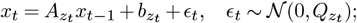

where each state *k* possesses its own set of rules (**A**_*k*_ for dynamics, **b**_*k*_ for bias, and **Q**_*k*_ for noise covariance) that describe the temporal progression of the low-dimensional patterns, accounting for Gaussian innovation noise *ϵ*_*t*_. Finally, the observed activity *y*_*t*_ is modeled as a linear projection of the continuous latent state *x*_*t*_ with noise:

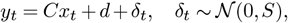

meaning that the BOLD signal is a linear combination of the latent patterns, adjusted by a bias vector *d*, and corrupted by observation noise δ_*t*_. Here, *C* ∈ ℝ^*N ×D*^ is the observation matrix, and *S* ∈ ℝ^*N ×N*^ is the covariance matrix of the observation noise. The complete set of parameters learned by the rSLDS model, which are optimized during inference using approximate variational inference methods (Linderman et al., 2017), is:

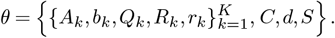

This comprehensive set includes parameters for the state transition dynamics, the distinct linear dynamical systems for each state, and the node-specific emission parameters that link the latent activity to the observed signals.

### K. Flow characteristic timescales

Thus, we calculated the flow on the top twenty HCS fractional occupancy participants balancing the dominant LCS fractional occupancy, which allowed us to quantify timescales of convergence across PC1 and PC2 (Fig. S13B) from the dynamic velocity (Nair et al., 2023).

The dynamic velocity quantified the speed of the activity in the state space, where the dynamics matrix *A*_*k*_ was computed. In specific, we separated it for the two dimensions (*D* = 2) and defined as 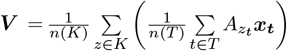, in consistency with (Nair et al., 2023); where *V*_1_ with *x*_1_ and *V*_2_ with *x*_2_ represent the dynamic velocities across PC1 and PC2 axes, respectively. Accordingly, we defined characteristic timescales as ***τ*** = 1*/****V***, which finally concluded to the timescales ratio between the two dimensions, *ϵ* = *τ*_*x*1_ */τ*_*x*2_ (Fig. S13).

### L. States transition

This switching mechanism is known as the Landauer’s principle (Landauer, 1961), it has been linked with information process and reliability of a device (Haken, 1977) and embodies the dynamics of the “labile brain” (Friston, 2000). On the other hand, the jump was identified as the case where the activity trajectory changed modes directly. The architecture of this mechanism has been studied extensively within the context of whole brain modeling of resting-state activity (Rabuffo et al., 2021; Fousek et al., 2024) through bistable neural masses, e.g. (Montbrió et al., 2015).

HMM was applied to the one-dimensional PC1 time series of the HCS states data, with two hidden states, to identify the two HCS modes corresponding to its bistable dynamics. Together with the LCS points a frequentist analysis was done, illustrated by Figure 4C, to judge for types of transitions. This led to the construction of a transition matrix, of shape 3×3, between LCS and the two HCS modes for each network. We constructed a simple periodic model that can account for three states (Fig. S15), namely

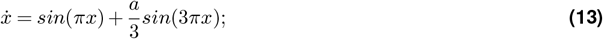

where *a* was the only free parameter that controlled for the number of solutions. For *a*_*th*_ ≃ 3.117, the model is close to bifurcation, since if *a > a*_*th*_ there are three solutions, else one. We simulated around a range of *a* values close to *a*_*th*_ and then applied simulation-based inference implemented via the sbi package (Tejero-Cantero et al., 2020).

## M. Acknowledgments

This project/research has received funding from the European Union’s Horizon Europe Programme under the Specific Grant Agreement No. 101147319 (EBRAINS 2.0 Project), No. 101137289 (Virtual Brain Twin Project), No. 101057429 (project environMENTAL), and government grant managed by the Agence Nationale de la Recherche reference ANR-22-PESN-0012 (France 2030 program). The funders had no role in study design, data collection and analysis, decision to publish, or preparation of the manuscript.

Data were provided [in part] by the Human Connectome Project, WU-Minn Consortium (Principal Investigators: David Van Essen and Kamil Ugurbil; 1U54MH091657) funded by the 16 NIH Institutes and Centers that support the NIH Blueprint for Neuroscience Research; and by the McDonnell Center for Systems Neuroscience at Washington University.

Data collection and sharing for this project was funded by the Alzheimer’s Disease Neuroimaging Initiative (ADNI) (National Institutes of Health Grant U01 AG024904) and DOD ADNI (Department of Defense award number W81XWH-12-2-0012). ADNI is funded by the National Institute on Aging, the National Institute of Biomedical Imaging and Bioengineering, and through generous contributions from the following: AbbVie, Alzheimer’s Association; Alzheimer’s Drug Discovery Foundation; Araclon Biotech; BioClinica, Inc.; Biogen; Bristol-Myers Squibb Company; CereSpir, Inc.; Cogstate; Eisai Inc.; Elan Pharmaceuticals, Inc.; Eli Lilly and Company; EuroImmun; F. Hoffmann-La Roche Ltd and its affiliated company Genentech, Inc.; Fujirebio; GE Healthcare; IXICO Ltd.; Janssen Alzheimer Immunotherapy Research & Development, LLC.; Johnson & Johnson Pharmaceutical Research & Development LLC.; Lumosity; Lundbeck; Merck & Co., Inc.; Meso Scale Diagnostics, LLC.; NeuroRx Research; Neurotrack Technologies; Novartis Pharmaceuticals Corporation; Pfizer Inc.; Piramal Imaging; Servier; Takeda Pharmaceutical Company; and Transition Therapeutics. The Canadian Institutes of Health Research is providing funds to support ADNI clinical sites in Canada. Private sector contributions are facilitated by the Foundation for the National Institutes of Health (www.fnih.org). The grantee organization is the Northern California Institute for Research and Education, and the study is coordinated by the Alzheimer’s Therapeutic Research Institute at the University of Southern California. ADNI data are disseminated by the Laboratory for Neuro Imaging at the University of Southern California.

## O. Information Sharing Statement

No new data were created or analyzed in this study. All code is available on GitHub.

## Supplementary Information

**Figure S1.**
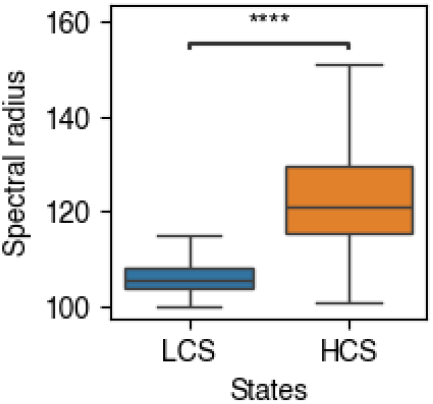
Reduced effective dimensionality during HCS compared to LCS. Significantly higher spectral radius, i.e. the first eigenvalue of the functional connectivity (FC) matrices, during HCS than LCS, demonstrates dimensionality reduction and thus more coordinative dynamics during the HCS.

**Figure S2.**
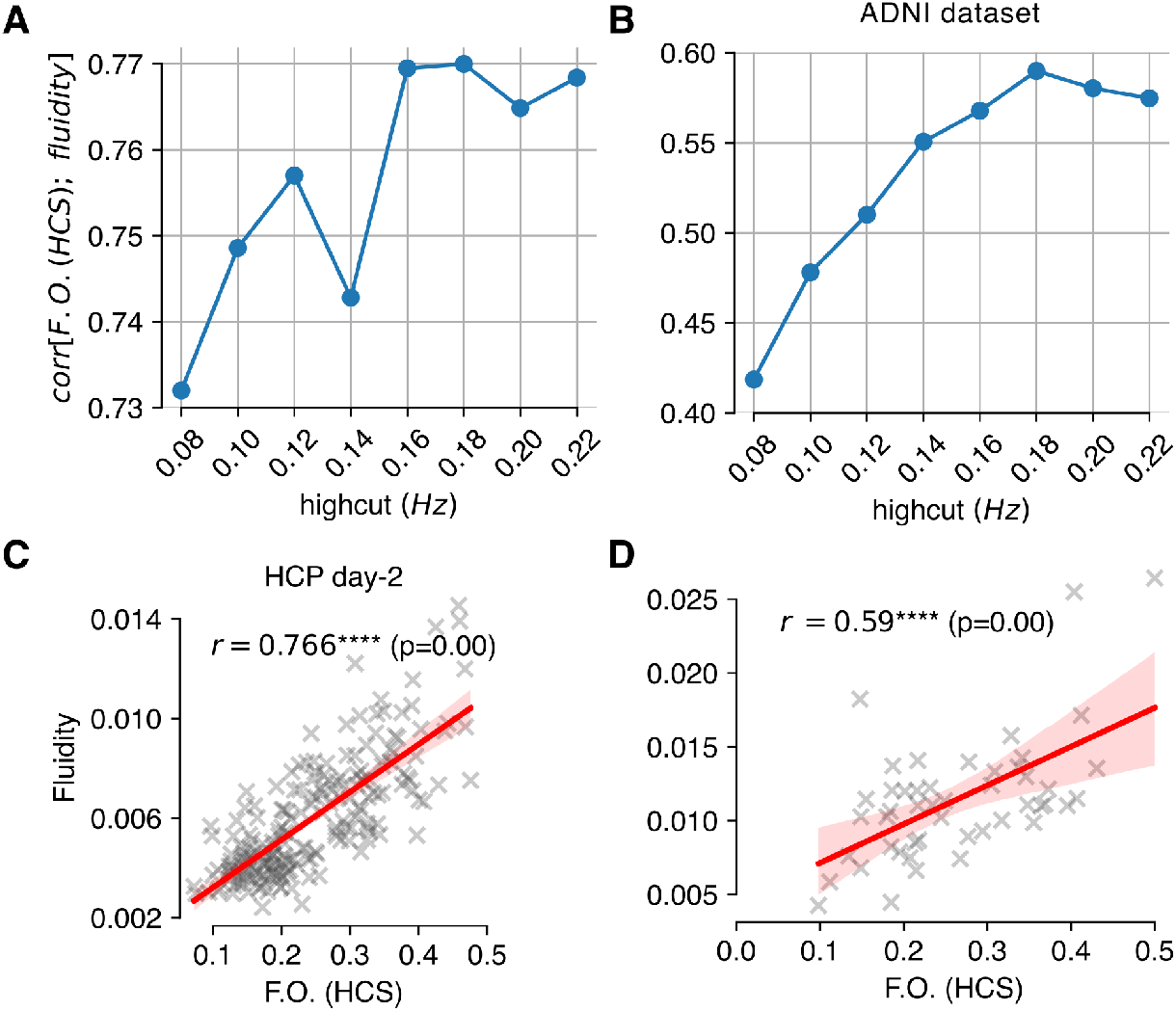
Fluidity metrics determine the bandpass highcut and show correlation with the fractional occupancy of the HCS for two high-resolution datasets. (**A**)The highcut at 0.18Hz yielded the best correlation of HCS fractional occupancy (FO) with fluidity, which was the value used to bandpass filter (lowcut=0.01Hz) the data. (**B**) Same result for the ADNI dataset. (**C**) Similar to Figure 1D, the fluidity values correlate (Pearson correlation) with the FO(HCS) across participants in the second-day HCP data (*r* = 0.766; *p* < 10^−4^). (**D**) Similar to (**C**), but for the ADNI dataset (*r* = 0.59; *p* < 10^−4^).

**Figure S3.**
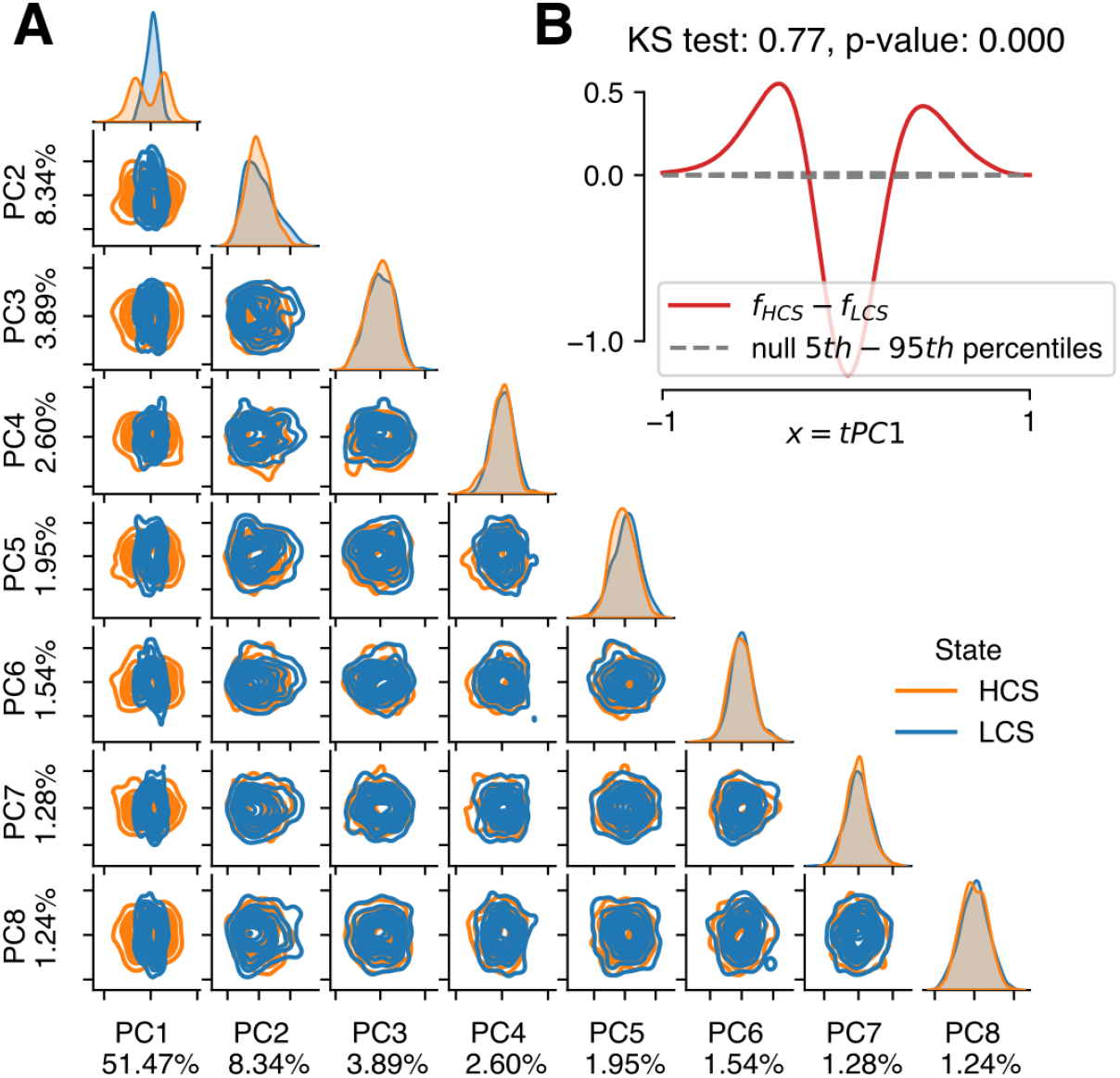
PCA across more components and quantification of significance. (**A**) Projections of the data from a test participant for both states on the first 8 PCs pairwise reveals that it is only the PC1 that distinguishes the LCS and HCS densities. (**B**) The difference of the one-dimensional estimated densities between LCS and HCS (red line) from the concatenated data of all participants significantly differs from the null distribution where the difference comes from LCS and HCS coming from the same distribution (grey dashed) (*KS* = 0.77; *p* < 1*e* − 5)

**Figure S4.**
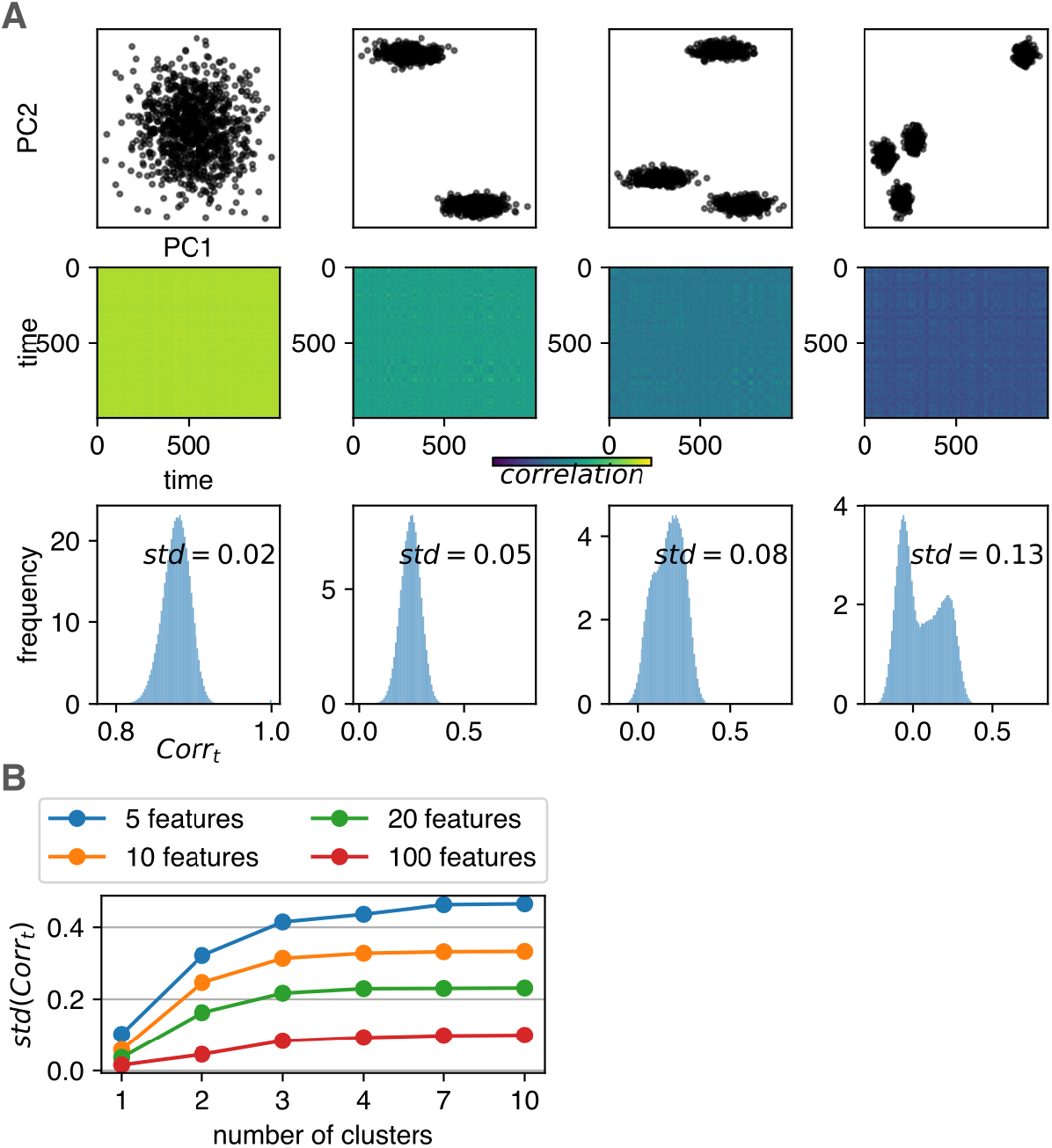
Sensitivity analysis for richness of coactivation patterns. (**A**) Timeseries that correspond to different number of clusters in a low-dimensional space yield correlations across time whose histograms acquire increasing standard deviations as the number of clusters increases. (**B**) This increase is stronger for data with less features (starting from 5), as the increasing trend weakens with a larger number of features, and plateaus faster.

**Figure S5.**
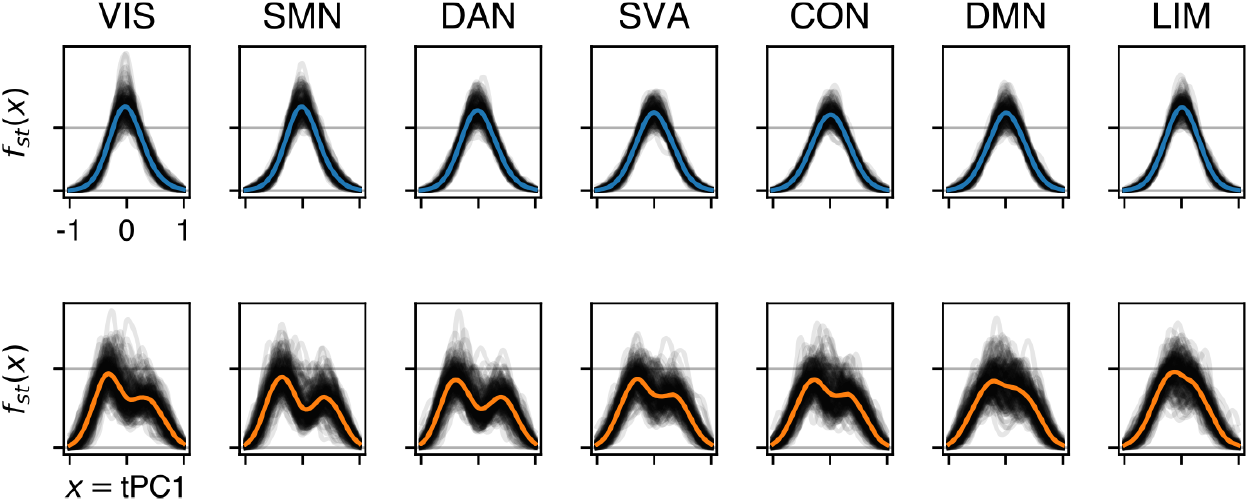
Observables stationary densities across RSNs for all participants. One-dimensional stationary densities *f*_*st*_(*x*), for *x* = *tPC*1, which is the PC1 scores, across the 7 RSNs, for the two different states, LCS and HCS, for all participants.

**Figure S6.**
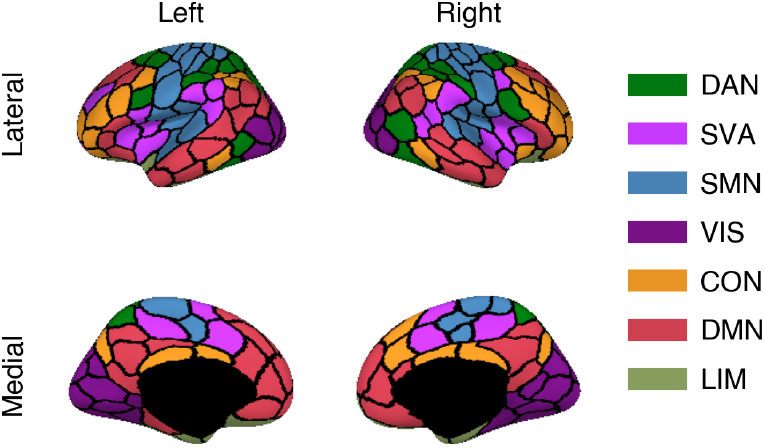
7 resting state networks. Anatomical map of the 7 resting state functional networks (RSNs) using the 200 regions of interest parcellation. These are the dorsal attention network (DAN), salient ventral attention (SVA) network, somato-motor network (SMN), visual (VIS) network, executive control network (CON), default mode network (DMN), and the limbic (LIM) network.

**Figure S7.**
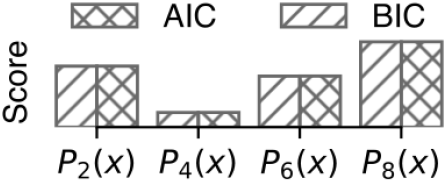
Model comparison. The normal forms, or *Thom’s unfoldings* (Methods E), of simpler (second-order *P*_2_(*x*)) and more complex (sixth-order *P*_6_(*x*), eighth order *P*_8_(*x*)) polynomials were fit to the data and compared to the fit for Eq. (6), here represented as *P*_4_(*x*). The Akaike and Bayesian information criteria quantified the trade-off between complexity and accuracy to demonstrate that *P*_4_(*x*) provides the best fit.

**Figure S8.**
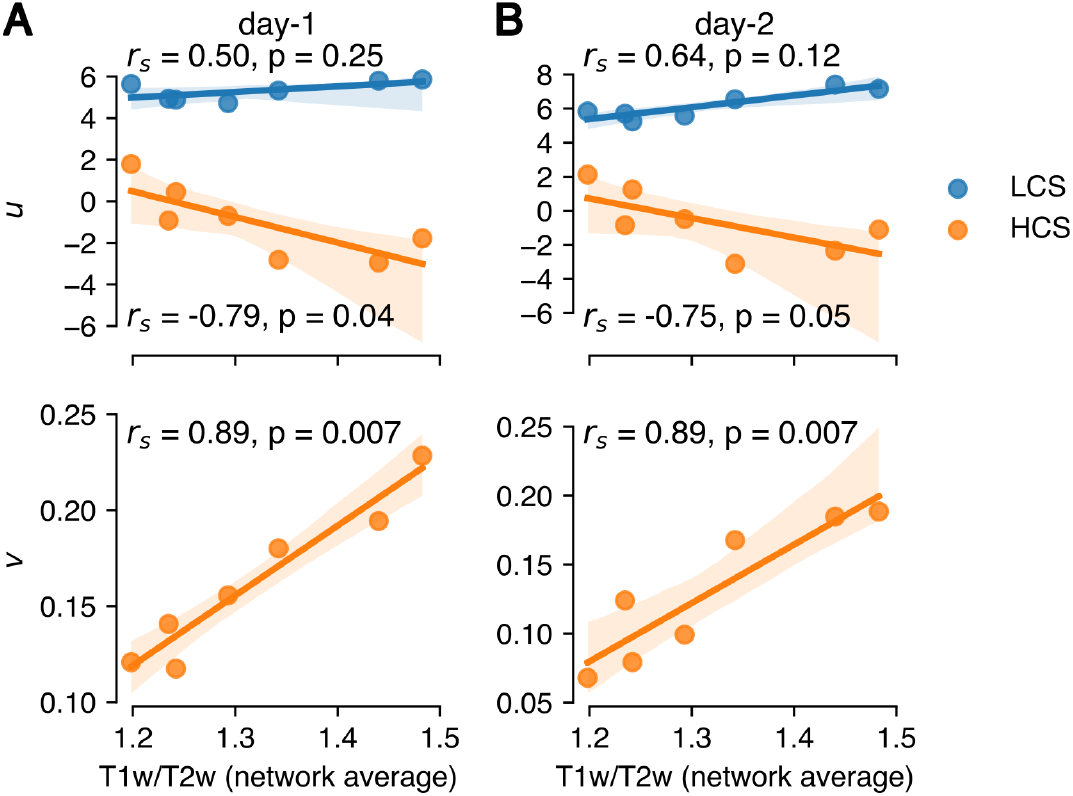
Parameters comparison with the average across subjects and networks T1w/T2w map. (**A**) Both *u* and *v* parameters show strong spatial correlations with the average T1w/T2w map (averaged across subjects and networks), with the HCS showing the most robust association. (**B**) Same for the HCP day-2 validation dataset.

**Figure S9.**
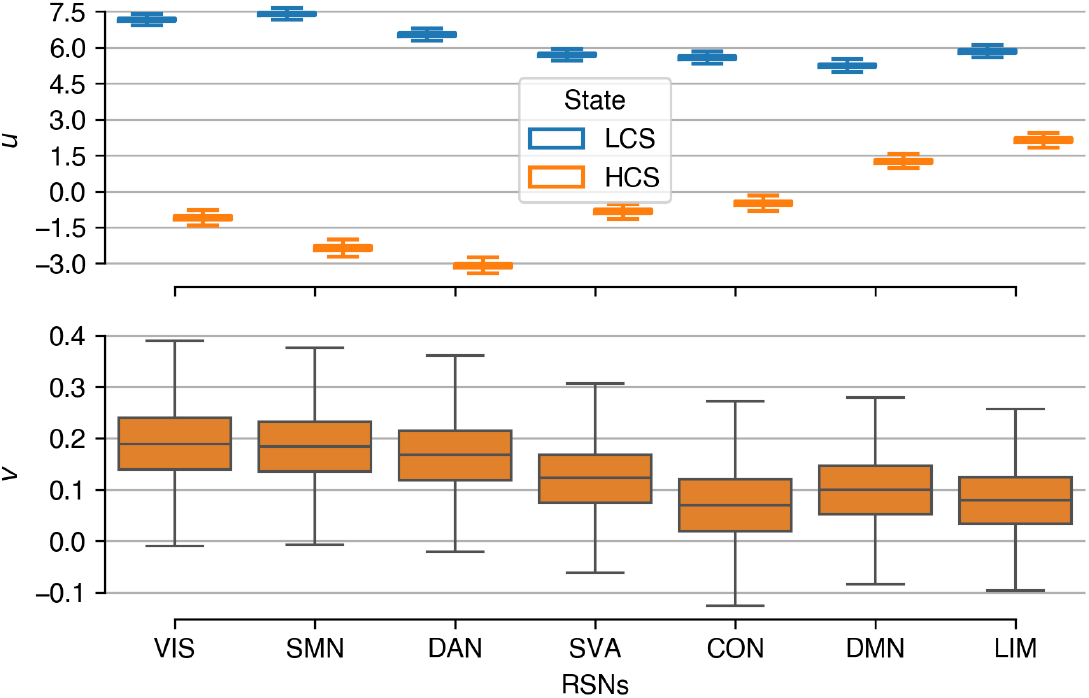
Second-day HCP data were analyzed and qualitatively replicated the original result. The *u* parameter reveals similar geometries for the RSNs as in the original result. The *v* parameter, again dynamically relevant only for HCS, showed even more pronounced trend following the functional gradient and acquired the highest value for the visual network as in the original result.

**Figure S10.**
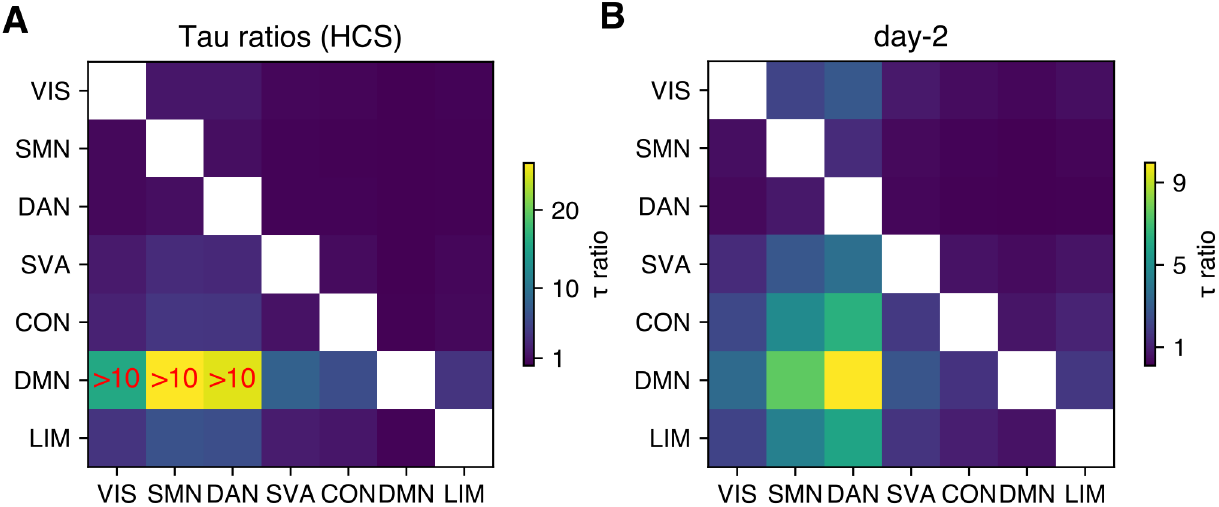
Timescale ratios between networks in HCS revealed that DMN acquired the slowest characteristic timescale. (**A**) The average timescale ratio (*τ*) was calculated pairwise across all RSNs and revealed that DMN operates in the slowest timescale, which even crossed the factor of 10 with periphery networks and DAN. Compared to SVA and other core networks it did not show such separation. (**B**) Similarly, the DMN acquired the slowest timescale, but did not cross the factor of 10 threshold for any of the networks.

**Figure S11.**
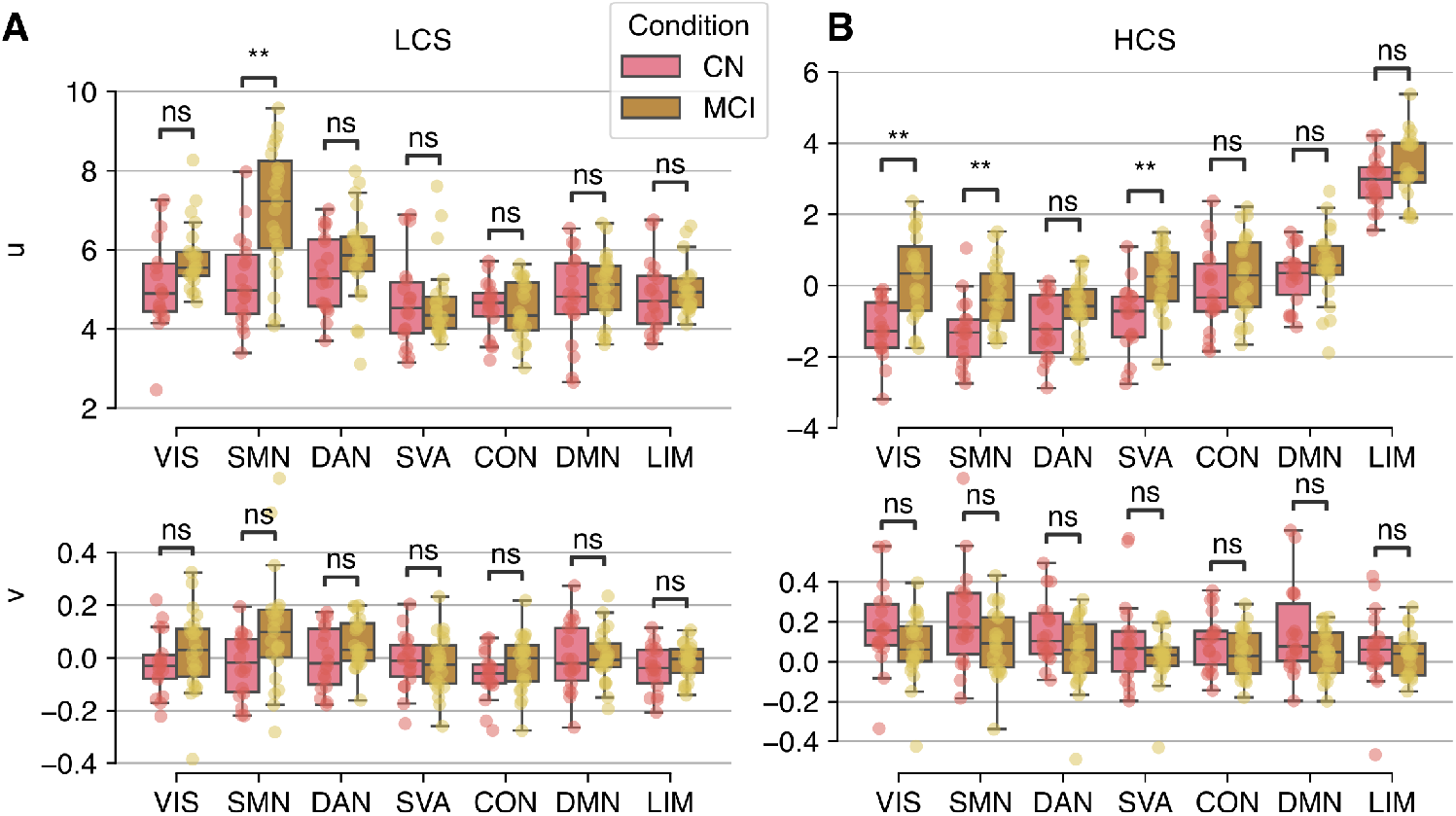
Subject-level comparisons of the inferred parameters *u* and *v* between control (CN) (P=18) and mildly cognitively impaired (MCI) (P=23) participants displays incohesive change of the parameters during LCS (significant difference only in SMN), but a consistent upwards drift for *u* of HCS across all RSNs and significantly for VIS, SMN and SVA. From all posterior samples across participants and networks for the inferred *u* and *v* parameters, only the average value for each participant is displayed for the two conditions, CN and MCI, for the two states. (**A**) In LCS, only the SMN had significantly higher *u* for the MCI, without a consistent drift across networks, either for *u* or *v*. (**B**) In HCS, all networks showed elevated *u* for MCI compared to CN, yet significant only for the peripheral networks (VIS and SMN) together with SVA (*p* < 10^−2^). Complementarily, the *v* parameter exhibited a consistent downwards drift, yet not significant (ns). Importantly, the CN distributions display similar organization as in Figure 3G.

**Figure S12.**
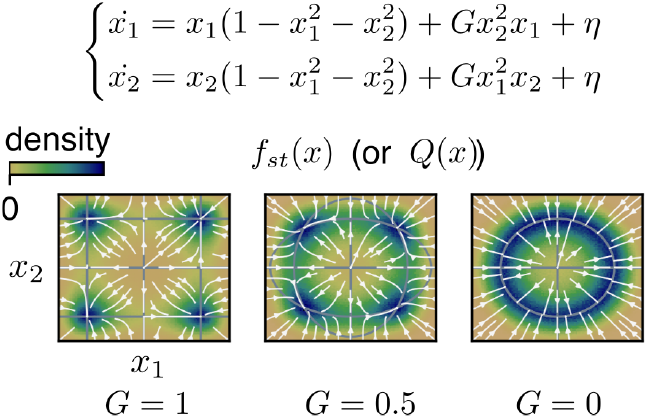
SFM generation from a bistable network. As the control parameter, *G*, breaks the symmetry (0<*G*<1) in this toy network of two nodes, *x*_1_ and *x*_2_, the SFM emerges along with timescales separation (*G* = 0.5). For *G* = 1, the nullclines form infinite straight lines; while for *G* = 0, the nullclines converge to a ring manifold.

**Figure S13.**
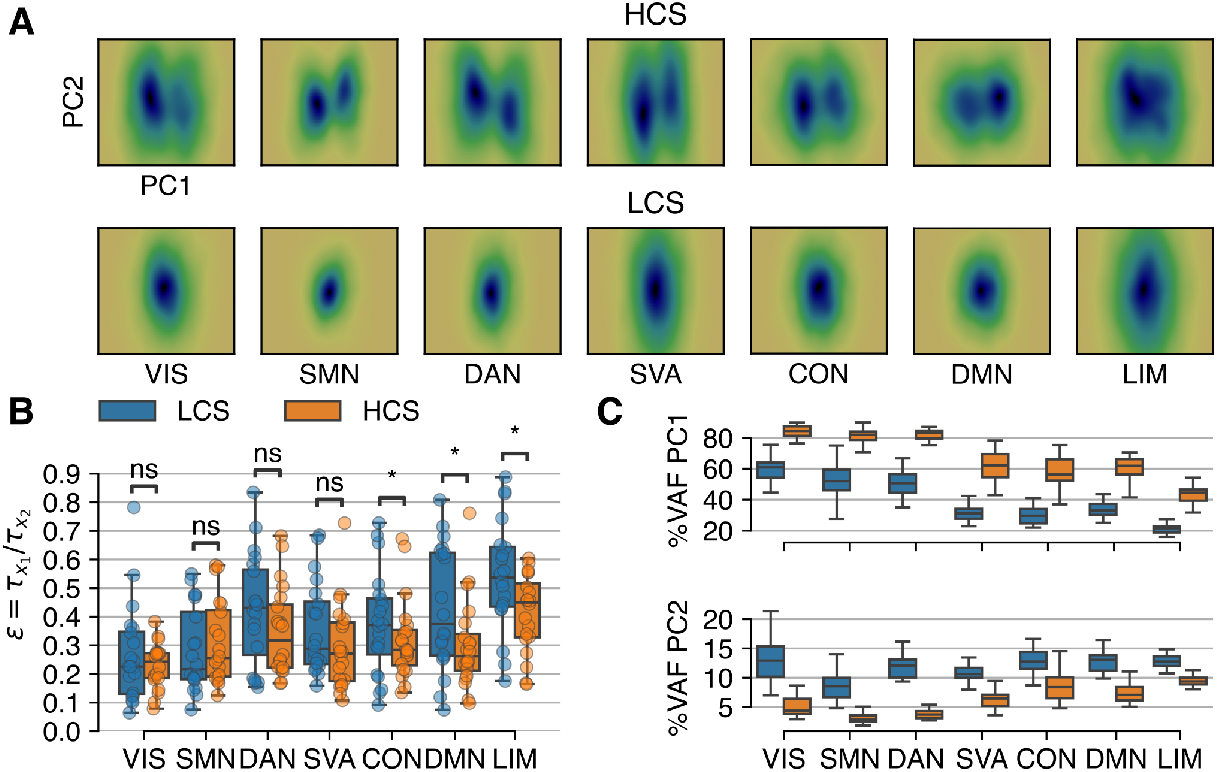
Attractor manifolds and timescales separation calculated for the 20 participants with the most balanced fractional occupancies between LCS and HCS, and thus best statistics. **A**Densities from the projections in the first two PCs from concatenated data of the 20 participants, for both HCS and LCS. Timescales separation 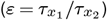 between *x*_1_ = *tPC*1 and *x*_2_ = *tPC*2 reveals that across *PC*2 the convergence to the stable fixed point is ~3 times slower (⟨*ε*⟩ ≃ 0.3). Consequently, the stable fixed point found from the previous PC1 analysis, converts into a slow manifold unfolding across PC2. This is true for both LCS and HCS. Notably, the core networks (LIM, CON and DMN) exhibited significantly slower convergence in HCS than LCS, which is consistent with their slower dynamics compared to the rest of RSNs, found from the PC1 analysis. (**C**) The observed timescales separation (*ε*), quantified by the *dynamic velocity* (Methods K), is not an artifact of the embedding, as the pattern of *ε* values across RSNs found in (**B**) is not reflected by the PC1, PC2 loadings.

**Figure S14.**
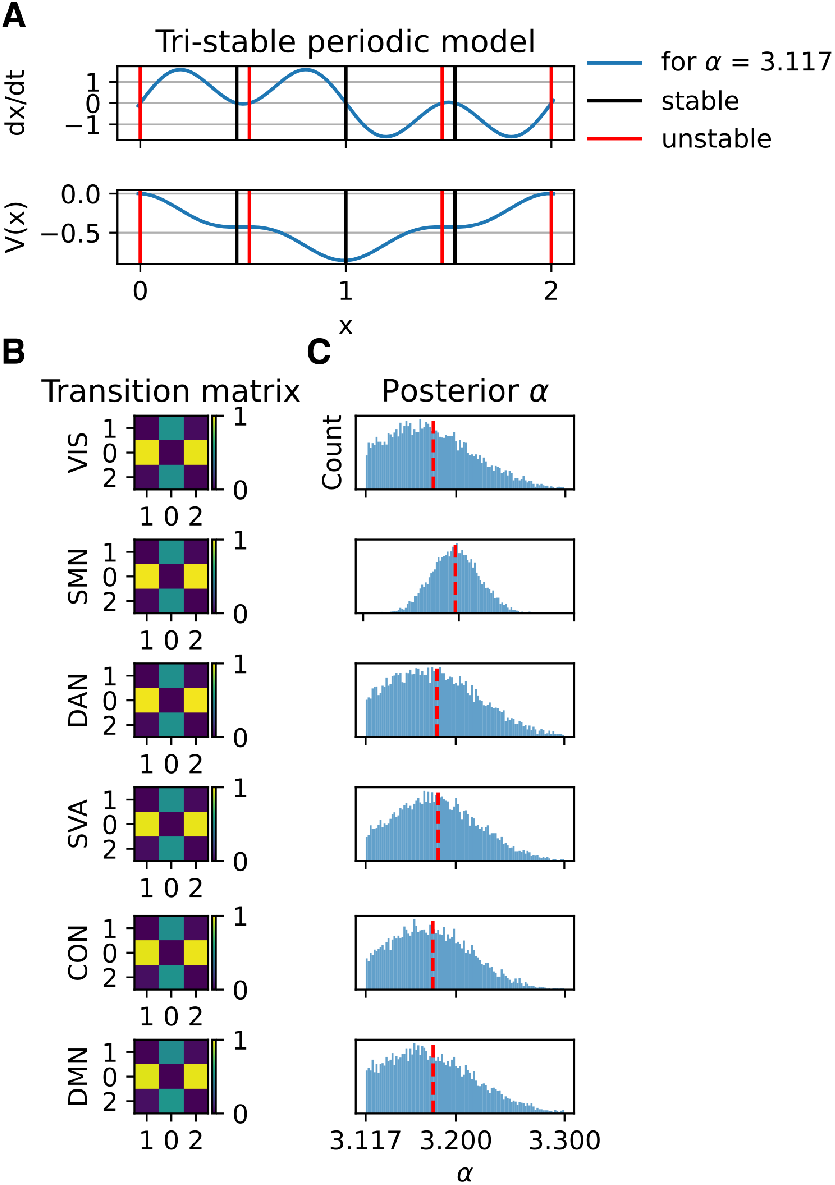
Qualitative sensitivity analysis based on the empirical transition matrices reveals that the system is metastable, namely close to a bifurcation point, where small changes can turn it from multistable to monostable. (**A**) A tri-stable periodic model, Eq. (13), with one bifurcation parameter (*α*) that controlled the existence of three or one fixed point, was adapted to capture the existence of three fixed points based on previous analysis across the PC1; that is one fixed point for the LCS and two for the HCS. (**B**) From the transitions found in the data between these three fixed points, transition matrices were formed for each RSN. (**C**) Using simulation-based inference, we inferred that *α* must be close to a bifurcation point, *α* ≲ 3.117, that transforms the tri-stable model to monostable, in order to support the values of the empirical transition matrices.

**Figure S15.**
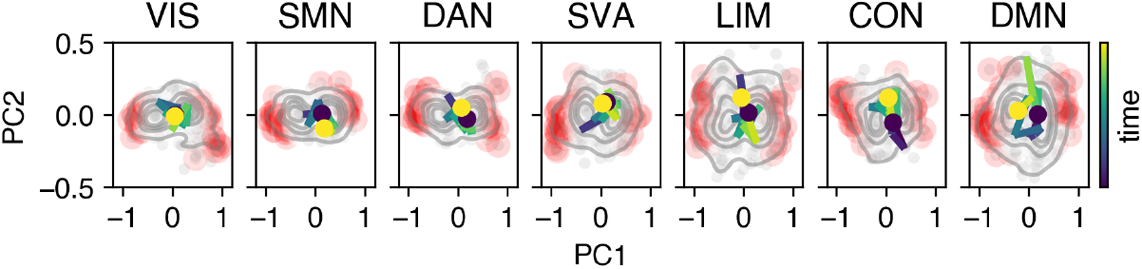
Exemplary *jump* without CAE takes place around 0. A jump that took place without the occurrence of a CAE is illustrated on the PC1-PC2 projection. The trajectory is around 0, suggesting a low-potential barrier at the time of the jump.

**Figure S16.**
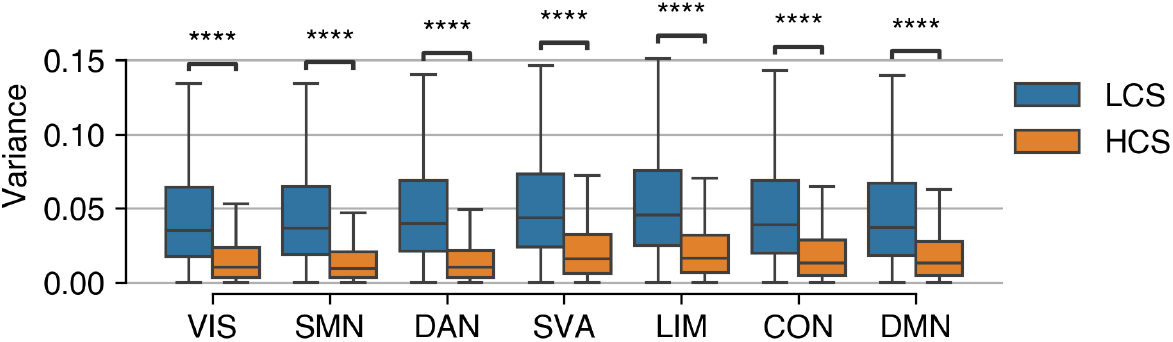
PC1 single trajectories variance in LCS is significantly larger than in HCS. Despite the larger variance of the stationary densities in HCS (Fig. S5), individual HCS trajectories are mainly locked into a single mode (rare jumps) and thus exhibit significantly smaller variance than LCS trajectories, measured from the projection onto the PC1.

**Figure S17.**
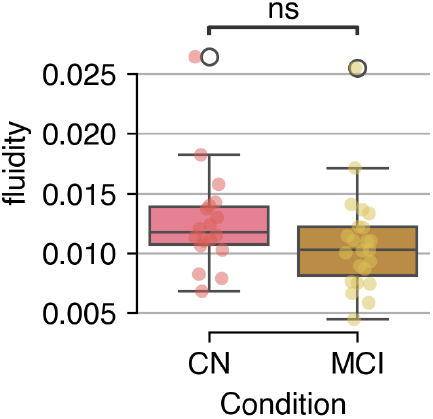
Fluidity values across participants cannot distinguish MCI from CN. Distributions of the fluidities across participants for CN and MCI showed no significant separation, with moderate effect size (*p* = 0.068, *d* = 0.69).

## Supplementary Tables

**Table S1.** Cohen’s *d* quantifies the effect size between CN and MCI. For the seven RSNs, *u*_*HCS*_ effect size is bigger than *u*_*LCS*_, except for SMN, where both effects are large. Moreover, for HCS, they clearly point to the same direction; namely, that *u*_*MCI*_ is larger than *u*_*CN*_. Same for *v*_*HCS*_, but now the effect size is smaller, and for SMN, DAN, LIM and CON, is smaller than *v*_*LCS*_. Effect sizes interpretation: 0.2 < *d* ≤ 0.5, small; 0.5 < *d* ≤ 0.8, moderate; 0.8 < *d* ≤ 1.3, large; *d* ≥ 1.3 very large.

|  | VIS | SMN | DAN | SVA | CON | DMN | LIM |
| --- | --- | --- | --- | --- | --- | --- | --- |
| $u_{LCS}$ | 0.88 | 1.46 | 0.51 | 0.08 | -0.01 | 0.33 | 0.35 |
| $u_{HCS}$ | 1.53 | 1.30 | 0.62 | 1.24 | 0.45 | 0.38 | 0.49 |
| $v_{LCS}$ | 0.44 | 0.81 | 0.65 | -0.03 | 0.53 | 0.15 | 0.45 |
| $v_{HCS}$ | -0.51 | -0.46 | -0.44 | -0.15 | -0.29 | -0.34 | -0.17 |

## Bibliography

Akaike, H., Petrov, B. N., and Csaki, F. Second international symposium on information theory, (1973).

Atasoy, S., Donnelly, I., and Pearson, J. (2016). Human brain networks function in connectome-specific harmonic waves. Nature communications, 7(1):10340.

Bennett, C. H. (1982). The thermodynamics of computation—a review. International Journal of Theoretical Physics, 21(12): 905–940.

Bingham, E., Chen, J. P., Jankowiak, M., Obermeyer, F., Pradhan, N., Karaletsos, T., Singh, R., Szerlip, P. A., Horsfall, P., and Goodman, N. D. (2019). Pyro: Deep universal probabilistic programming. J. Mach. Learn. Res., 20:28:1–28:6.

Bishop, C. M. Pattern Recognition and Machine Learning (Information Science and Statistics). Springer-Verlag, Berlin, Heidelberg, (2006). ISBN 0387310738.

Biswal, B., Zerrin Yetkin, F., Haughton, V. M., and Hyde, J. S. (1995). Functional connectivity in the motor cortex of resting human brain using echo-planar mri. Magnetic resonance in medicine, 34(4):537–541.

Bouzigues, A., Godefroy, V., Le Du, V., Russell, L. L., Houot, M., Le Ber, I., Batrancourt, B., Levy, R., Warren, J. D., Rohrer, J. D., Margulies, D. S., and Migliaccio, R. (2024). Disruption of macroscale functional network organisation in patients with frontotemporal dementia. Molecular Psychiatry, pages 1–12. doi: 10.1038/s41380-024-02847-4.

Breyton, M., Fousek, J., Rabuffo, G., Sorrentino, P., Kusch, L., Massimini, M., Petkoski, S., and Jirsa, V. Large-scale brain signatures of fluid dynamics and responsiveness linked to consciousness. Preprint, Neuroscience, (2023).

Buzsáki, G. and Draguhn, A. (2004). Neuronal Oscillations in Cortical Networks. Science, 304(5679):1926–1929. doi: 10.1126/science.1099745.

Cabral, J., Vidaurre, D., Marques, P., Magalhães, R., Silva Moreira, P., Miguel Soares, J., Deco, G., Sousa, N., and Kringelbach, M. L. (2017). Cognitive performance in healthy older adults relates to spontaneous switching between states of functional connectivity during rest. Scientific Reports, 7(1):5135. doi: 10.1038/s41598-017-05425-7.

Christoff, K., Irving, Z. C., Fox, K. C., Spreng, R. N., and Andrews-Hanna, J. R. (2016). Mind-wandering as spontaneous thought: a dynamic framework. Nature reviews neuroscience, 17(11):718–731.

Clawson, W., Vicente, A. F., Ferraris, M., Bernard, C., Battaglia, D., and Quilichini, P. P. (2019). Computing hubs in the hippocampus and cortex. Science Advances, 5(6):eaax4843. doi: 10.1126/sciadv.aax4843.

Crauel, H. and Flandoli, F. (1994). Attractors for random dynamical systems. Probability Theory and Related Fields, 100(3): 365–393.

Damasio, A. R. and Damasio, H. (1994). Cortical systems for retrieval of concrete knowledge: The convergence zone framework.

Deco, G., Jirsa, V. K., and McIntosh, A. R. (2011). Emerging concepts for the dynamical organization of resting-state activity in the brain. Nature Reviews Neuroscience, 12(1):43–56. doi: 10.1038/nrn2961.

Deco, G., Senden, M., and Jirsa, V. (2012). How anatomy shapes dynamics: a semi-analytical study of the brain at rest by a simple spin model. Frontiers in computational neuroscience, 6:68.

Dehaene, S. and Cohen, L. (2007). Cultural Recycling of Cortical Maps. Neuron, 56(2):384–398. doi: 10.1016/j.neuron.2007.10.004.

Dimakou, A., Pezzulo, G., Zangrossi, A., and Corbetta, M. (2025). The predictive nature of spontaneous brain activity across scales and species. Neuron, 113(9):1310–1332. doi: 10.1016/j.neuron.2025.02.009.

Domhof, J., Eickhoff, S., Jung, K., and Popovych, O. Parcellation-based structural and resting-state functional brain connectomes of a healthy cohort. Technical report, Gehirn & Verhalten, (2021).

Englert, R., Kincses, B., Kotikalapudi, R., Gallitto, G., Li, J., Hoffschlag, K., Woo, C.-W., Wager, T. D., Timmann, D., Bingel, U., and Spisak, T. (2024). Connectome-Based Attractor Dynamics Underlie Brain Activity in Rest, Task, and Disease. eLife, 13. doi: 10.7554/eLife.98725.1.

Esfahlani, F., Jo, Y., Faskowitz, J., Byrge, L., Kennedy, D. P., Sporns, O., and Betzel, R. F. (2020). High-amplitude cofluctuations in cortical activity drive functional connectivity. Proceedings of the National Academy of Sciences, 117(45):28393–28401. doi: 10.1073/pnas.2005531117.

Fousek, J., Rabuffo, G., Gudibanda, K., Sheheitli, H., Petkoski, S., and Jirsa, V. (2024). Symmetry breaking organizes the brain’s resting state manifold. Scientific Reports, 14(1):31970. doi: 10.1038/s41598-024-83542-w.

Friston, K. (2013). Life as we know it. Journal of the Royal Society Interface, 10(86):20130475.

Friston, K., Da Costa, L., Sajid, N., Heins, C., Ueltzhöffer, K., Pavliotis, G. A., and Parr, T. (2023). The free energy principle made simpler but not too simple. Physics Reports, 1024:1–29.

Friston, K. J. (2000). The labile brain. i. neuronal transients and nonlinear coupling. Philosophical Transactions of the Royal Society of London. Series B: Biological Sciences, 355(1394):215–236.

Ghosh, A., Rho, Y., McIntosh, A. R., Kötter, R., and Jirsa, V. K. (2008). Noise during rest enables the exploration of the brain’s dynamic repertoire. PLoS computational biology, 4(10):e1000196.

Glasser, M. F., Sotiropoulos, S. N., Wilson, J. A., Coalson, T. S., Fischl, B., Andersson, J. L., Xu, J., Jbabdi, S., Webster, M., Polimeni, J. R., et al. (2013). The minimal preprocessing pipelines for the human connectome project. Neuroimage, 80: 105–124. doi: 10.1016/j.humov.2008.02.015.Changes.

Golos, M., Jirsa, V., and Daucé, E. (2015). Multistability in large scale models of brain activity. PLoS computational biology, 11 (12):e1004644.

Gonzalez-Castillo, J., Hoy, C. W., Handwerker, D. A., Robinson, M. E., Buchanan, L. C., Saad, Z. S., and Bandettini, P. A. (2015). Tracking ongoing cognition in individuals using brief, whole-brain functional connectivity patterns. Proceedings of the National Academy of Sciences, 112(28):8762–8767.

Haken, H. (1977). Synergetics. Physics Bulletin, 28(9):412.

Hancock, F., Rosas, F. E., Luppi, A. I., Zhang, M., Mediano, P. A. M., Cabral, J., Deco, G., Kringelbach, M. L., Breakspear, M., Kelso, J. A. S., and Turkheimer, F. E. (2024). Metastability demystified — the foundational past, the pragmatic present and the promising future. Nature Reviews Neuroscience, pages 1–19. doi: 10.1038/s41583-024-00883-1.

Hansen, E. C., Battaglia, D., Spiegler, A., Deco, G., and Jirsa, V. K. (2015). Functional connectivity dynamics: Modeling the switching behavior of the resting state. NeuroImage, 105:525–535. doi: 10.1016/j.neuroimage.2014.11.001.

Hashemi, M., Depannemaecker, D., Saggio, M., Triebkorn, P., Rabuffo, G., Fousek, J., Ziaeemehr, A., Sip, V., Athanasiadis, A., Breyton, M., Woodman, M., Wang, H., Petkoski, S., Sorrentino, P., and Jirsa, V. (2025). Principles and operation of virtual brain twins. IEEE Reviews in Biomedical Engineering, pages 1–29. doi: 10.1109/RBME.2025.3562951.

He, B. J. (2013). Spontaneous and task-evoked brain activity negatively interact. Journal of Neuroscience, 33(11):4672–4682.

Honey, C. J., Kötter, R., Breakspear, M., and Sporns, O. (2007). Network structure of cerebral cortex shapes functional connectivity on multiple time scales. Proceedings of the National Academy of Sciences, 104(24):10240–10245.

Honey, C. J., Thesen, T., Donner, T. H., Silbert, L. J., Carlson, C. E., Devinsky, O., Doyle, W. K., Rubin, N., Heeger, D. J., and Hasson, U. (2012). Slow cortical dynamics and the accumulation of information over long timescales. Neuron, 76(2):423–434.

Hutchison, R. M., Womelsdorf, T., Allen, E. A., Bandettini, P. A., Calhoun, V. D., Corbetta, M., Della Penna, S., Duyn, J. H., Glover, G. H., Gonzalez-Castillo, J., et al. (2013). Dynamic functional connectivity: promise, issues, and interpretations. Neuroimage, 80:360–378.

Huys, R., Perdikis, D., and Jirsa, V. K. (2014). Functional architectures and structured flows on manifolds: a dynamical framework for motor behavior. Psychological review, 121(3):302.

Iraji, A., Faghiri, A., Fu, Z., Kochunov, P., Adhikari, B., Belger, A., Ford, J., McEwen, S., Mathalon, D., Pearlson, G., Potkin, S., Preda, A., Turner, J., Van Erp, T., Chang, C., and Calhoun, V. (2022). Moving beyond the ‘CAP’ of the Iceberg: Intrinsic connectivity networks in fMRI are continuously engaging and overlapping. NeuroImage, 251:119013. doi: 10.1016/j.neuroimage.2022.119013.

Ito, T., Hearne, L. J., and Cole, M. W. (2020). A cortical hierarchy of localized and distributed processes revealed via dissociation of task activations, connectivity changes, and intrinsic timescales. NeuroImage, 221:117141. doi: 10.1016/j.neuroimage.2020.117141.

Jaynes, E. T. (1957). Information theory and statistical mechanics. Physical review, 106(4):620.

Jaynes, E. T. (2007). Prior probabilities. IEEE Transactions on systems science and cybernetics, 4(3):227–241.

Jirsa, V. and Sheheitli, H. (2022). Entropy, free energy, symmetry and dynamics in the brain. Journal of Physics: Complexity, 3 (1):015007. doi: 10.1088/2632-072X/ac4bec.

Kelso, J. A. S. Dynamic Patterns: The Self-organization of Brain and Behavior. MIT Press, (1995). ISBN 978-0-262-61131-2.

Khona, M. and Fiete, I. R. (2022). Attractor and integrator networks in the brain. Nature Reviews Neuroscience, 23(12):744–766. doi: 10.1038/s41583-022-00642-0.

Kiebel, S. J., Daunizeau, J., and Friston, K. J. (2008). A hierarchy of time-scales and the brain. PLoS computational biology, 4 (11):e1000209.

Kong, X., Kong, R., Orban, C., Wang, P., Zhang, S., Anderson, K., Holmes, A., Murray, J. D., Deco, G., Van Den Heuvel, M., and Yeo, B. T. T. (2021). Sensory-motor cortices shape functional connectivity dynamics in the human brain. Nature Communications, 12(1):6373. doi: 10.1038/s41467-021-26704-y.

Kringelbach, M. L., Cruzat, J., Cabral, J., Knudsen, G. M., Carhart-Harris, R., Whybrow, P. C., Logothetis, N. K., and Deco, G. (2020). Dynamic coupling of whole-brain neuronal and neurotransmitter systems. Proceedings of the National Academy of Sciences, 117(17):9566–9576. doi: 10.1073/pnas.1921475117.

Lakatos, P., Gross, J., and Thut, G. (2019). A new unifying account of the roles of neuronal entrainment. Current Biology, 29(18): R890–R905.

Landauer, R. (1961). Irreversibility and heat generation in the computing process. IBM journal of research and development, 5 (3):183–191.

Lavanga, M., Stumme, J., Yalcinkaya, B. H., Fousek, J., Jockwitz, C., Sheheitli, H., Bittner, N., Hashemi, M., Petkoski, S., Caspers, S., and Jirsa, V. (2023). The virtual aging brain: Causal inference supports interhemispheric dedifferentiation in healthy aging. NeuroImage, 283:120403. doi: 10.1016/j.neuroimage.2023.120403.

Linderman, S., Johnson, M., Miller, A., Adams, R., Blei, D., and Paninski, L. Bayesian learning and inference in recurrent switching linear dynamical systems. in Artificial intelligence and statistics, pages 914–922. PMLR, (2017).

Liu, X. and Duyn, J. H. (2013). Time-varying functional network information extracted from brief instances of spontaneous brain activity. Proceedings of the National Academy of Sciences, 110(11):4392–4397. doi: 10.1073/pnas.1216856110.

Margulies, D. S., Ghosh, S. S., Goulas, A., Falkiewicz, M., Huntenburg, J. M., Langs, G., Bezgin, G., Eickhoff, S. B., Castellanos, F. X., Petrides, M., Jefferies, E., and Smallwood, J. (2016). Situating the default-mode network along a principal gradient of macroscale cortical organization. Proceedings of the National Academy of Sciences, 113(44):12574–12579. doi: 10.1073/pnas.1608282113.

Mazzara, C., Gambino, G., Duma, G. M., Neri, M., Giglia, G., Piccoli, T., Guajana, F., Comelli, A., Tuttolomondo, A., Migliore, M., and Sorrentino, P. (2025). Brain fluidity as a functional marker of tau-related neurodegeneration in alzheimer’s disease. doi: 10.1101/2025.07.04.25330744.

Montbrió, E., Pazó, D., and Roxin, A. (2015). Macroscopic Description for Networks of Spiking Neurons. Physical Review X, 5 (2):021028. doi: 10.1103/PhysRevX.5.021028.

Monto, S., Palva, S., Voipio, J., and Palva, J. M. (2008). Very slow eeg fluctuations predict the dynamics of stimulus detection and oscillation amplitudes in humans. Journal of Neuroscience, 28(33):8268–8272.

Murray, J. D., Bernacchia, A., Freedman, D. J., Romo, R., Wallis, J. D., Cai, X., Padoa-Schioppa, C., Pasternak, T., Seo, H., Lee, D., and Wang, X.-J. (2014). A hierarchy of intrinsic timescales across primate cortex. Nature Neuroscience, 17(12): 1661–1663. doi: 10.1038/nn.3862.

Naik, S., Banerjee, A., Bapi, R. S., Deco, G., and Roy, D. (2017). Metastability in senescence. Trends in cognitive sciences, 21 (7):509–521.

Nair, A., Karigo, T., Yang, B., Ganguli, S., Schnitzer, M. J., Linderman, S. W., Anderson, D. J., and Kennedy, A. (2023). An approximate line attractor in the hypothalamus encodes an aggressive state. Cell, 186(1):178–193.e15. doi: 10.1016/j.cell.2022.11.027.

Owens, C. D., Pinto, C. B., Mukli, P., Gulej, R., Velez, F. S., Detwiler, S., Olay, L., Hoffmeister, J. R., Szarvas, Z., Muranyi, M., et al. (2024). Neurovascular coupling, functional connectivity, and cerebrovascular endothelial extracellular vesicles as biomarkers of mild cognitive impairment. Alzheimer’s & Dementia, 20(8):5590–5606.

Pang, J. C., Aquino, K. M., Oldehinkel, M., Robinson, P. A., Fulcher, B. D., Breakspear, M., and Fornito, A. (2023). Geometric constraints on human brain function. Nature, 618(7965):566–574. doi: 10.1038/s41586-023-06098-1.

Parr, T., Pezzulo, G., and Friston, K. J. Active Inference: The Free Energy Principle in Mind, Brain, and Behavior. The MIT Press. The MIT Press, Cambridge, (2022). ISBN 978-0-262-04535-3 978-0-262-36997-8.

Peng, X., Liu, Q., Hubbard, C. S., Wang, D., Zhu, W., Fox, M. D., and Liu, H. (2023). Robust dynamic brain coactivation states estimated in individuals. Science Advances, 9(3):eabq8566. doi: 10.1126/sciadv.abq8566.

Perdikis, D., Huys, R., and Jirsa, V. K. (2011). Time Scale Hierarchies in the Functional Organization of Complex Behaviors. PLoS Computational Biology, 7(9):e1002198. doi: 10.1371/journal.pcbi.1002198.

Petersen, S. E. and Sporns, O. (2015). Brain Networks and Cognitive Architectures. Neuron, 88(1):207–219. doi: 10.1016/j.neuron.2015.09.027.

Phan, D., Pradhan, N., and Jankowiak, M. (2019). Composable effects for flexible and accelerated probabilistic programming in numpyro. arXiv preprint arXiv:1912.11554.

Pillai, A. S. and Jirsa, V. K. (2017). Symmetry Breaking in Space-Time Hierarchies Shapes Brain Dynamics and Behavior. Neuron, 94(5):1010–1026. doi: 10.1016/j.neuron.2017.05.013.

Ponce-Alvarez, A., He, B. J., Hagmann, P., and Deco, G. (2015). Task-driven activity reduces the cortical activity space of the brain: experiment and whole-brain modeling. PLoS computational biology, 11(8):e1004445.

Power, J. D., Cohen, A. L., Nelson, S. M., Wig, G. S., Barnes, K. A., Church, J. A., Vogel, A. C., Laumann, T. O., Miezin, F. M., Schlaggar, B. L., et al. (2011). Functional network organization of the human brain. Neuron, 72(4):665–678.

Rabuffo, G., Fousek, J., Bernard, C., and Jirsa, V. (2021). Neuronal Cascades Shape Whole-Brain Functional Dynamics at Rest. eNeuro, 8(5). doi: 10.1523/ENEURO.0283-21.2021.

Rabuffo, G., Lokossou, H.-A., Li, Z., Ziaee-Mehr, A., Hashemi, M., Quilichini, P. P., Ghestem, A., Arab, O., Esclapez, M., Verma, P., et al. (2025). Mapping global brain reconfigurations following local targeted manipulations. Proceedings of the National Academy of Sciences, 122(16):e2405706122.

Raichle, M. E. (2011). The Restless Brain. 1(1):3–12. doi: 10.1089/brain.2011.0019.

Raut, R. V., Snyder, A. Z., Mitra, A., Yellin, D., Fujii, N., Malach, R., and Raichle, M. E. (2021). Global waves synchronize the brain’s functional systems with fluctuating arousal. Science advances, 7(30):eabf2709.

Risken, H. Fokker-planck equation. in The Fokker-Planck equation: methods of solution and applications, pages 63–95. Springer, (1989).

Ságodi, Á., Martín-Sánchez, G., Sokol, P., and Park, M. (2024). Back to the continuous attractor. Advances in Neural Information Processing Systems, 37:66856–66906.

Salimi-Khorshidi, G., Douaud, G., Beckmann, C. F., Glasser, M. F., Griffanti, L., and Smith, S. M. (2014). Automatic denoising of functional mri data: combining independent component analysis and hierarchical fusion of classifiers. Neuroimage, 90: 449–468. doi: 10.1016/j.neuroimage.2013.11.046.

Salthe, S. N. Evolving hierarchical systems: their structure and representation. Columbia University Press, (1985).

Schaefer, A., Kong, R., Gordon, E. M., Laumann, T. O., Zuo, X.-N., Holmes, A. J., Eickhoff, S. B., and Yeo, B. T. T. (2018). Local-Global Parcellation of the Human Cerebral Cortex from Intrinsic Functional Connectivity MRI. Cerebral Cortex, 28(9): 3095–3114. doi: 10.1093/cercor/bhx179.

Schneider, S., Lee, J. H., and Mathis, M. W. (2023). Learnable latent embeddings for joint behavioural and neural analysis. 617 (7960):360–368. doi: 10.1038/s41586-023-06031-6.

Schöner, G. and Kelso, J. S. (1988). Dynamic pattern generation in behavioral and neural systems. Science, 239(4847): 1513–1520.

Schroeder, C. E. and Lakatos, P. (2009). Low-frequency neuronal oscillations as instruments of sensory selection. Trends in neurosciences, 32(1):9–18.

Schwarz, G. (1978). Estimating the dimension of a model. The annals of statistics, pages 461–464.

Senden, M., Reuter, N., van den Heuvel, M. P., Goebel, R., and Deco, G. (2017). Cortical rich club regions can organize state-dependent functional network formation by engaging in oscillatory behavior. NeuroImage, 146:561–574.

Shine, J. M., Breakspear, M., Bell, P. T., Ehgoetz Martens, K. A., Shine, R., Koyejo, O., Sporns, O., and Poldrack, R. A. (2019). Human cognition involves the dynamic integration of neural activity and neuromodulatory systems. Nature Neuroscience, 22 (2):289–296. doi: 10.1038/s41593-018-0312-0.

Smallwood, J., Bernhardt, B. C., Leech, R., Bzdok, D., Jefferies, E., and Margulies, D. S. (2021). The default mode network in cognition: A topographical perspective. Nature Reviews Neuroscience, 22(8):503–513. doi: 10.1038/s41583-021-00474-4.

Smith, S. M., Beckmann, C. F., Andersson, J., Auerbach, E. J., Bijsterbosch, J., Douaud, G., Duff, E., Feinberg, D. A., Griffanti, L., Harms, M. P., et al. (2013). Resting-state fmri in the human connectome project. Neuroimage, 80:144–168.

Stratonovich, R. L. Topics in the theory of random noise, volume 2. CRC Press, (1967).

Strogatz, S. H. Nonlinear dynamics and chaos: with applications to physics, biology, chemistry, and engineering. Chapman and Hall/CRC, (2024).

Tejero-Cantero, A., Boelts, J., Deistler, M., Lueckmann, J.-M., Durkan, C., Gonçalves, P. J., Greenberg, D. S., and Macke, J. H. (2020). sbi: A toolkit for simulation-based inference. Journal of Open Source Software, 5(52):2505. doi: 10.21105/joss.02505.

Thomas Yeo, B., Krienen, F. M., Sepulcre, J., Sabuncu, M. R., Lashkari, D., Hollinshead, M., Roffman, J. L., Smoller, J. W., Zöllei, L., Polimeni, J. R., et al. (2011). The organization of the human cerebral cortex estimated by intrinsic functional connectivity. Journal of neurophysiology, 106(3):1125–1165.

Van Essen, D. C., Ugurbil, K., Auerbach, E., Barch, D., Behrens, T. E., Bucholz, R., Chang, A., Chen, L., Corbetta, M., Curtiss, S. W., et al. (2012). The human connectome project: a data acquisition perspective. Neuroimage, 62(4):2222–2231.

Vidaurre, D., Smith, S. M., and Woolrich, M. W. (2017). Brain network dynamics are hierarchically organized in time. Proceedings of the National Academy of Sciences, 114(48):12827–12832. doi: 10.1073/pnas.1705120114.

Vinograd, A., Nair, A., Kim, J. H., Linderman, S. W., and Anderson, D. J. (2024). Causal evidence of a line attractor encoding an affective state. Nature, 634(8035):910–918.

Vohryzek, J., Sanz-Perl, Y., Kringelbach, M. L., and Deco, G. (2025). Human brain dynamics are shaped by rare long-range connections over and above cortical geometry. Proceedings of the National Academy of Sciences, 122(1):e2415102122.

Woodman, M. M. and Jirsa, V. K. (2013). Emergent Dynamics from Spiking Neuron Networks through Symmetry Breaking of Connectivity. PLoS ONE, 8(5):e64339. doi: 10.1371/journal.pone.0064339.

Wu, Y.-h., Podvalny, E., Levinson, M., and He, B. J. (2024). Network mechanisms of ongoing brain activity’s influence on conscious visual perception. Nature communications, 15(1):5720.

Zalesky, A., Fornito, A., Cocchi, L., Gollo, L. L., and Breakspear, M. (2014). Time-resolved resting-state brain networks. Proceedings of the National Academy of Sciences, 111(28):10341–10346.

Zeeman, E. C. Catastrophe theory. in Structural Stability in Physics: Proceedings of Two International Symposia on Applications of Catastrophe Theory and Topological Concepts in Physics Tübingen, Fed. Rep. of Germany, May 2–6 and December 11–14, 1978, pages 12–22. Springer, (1979).

